# Substrate-Specific Effects of Natural Genetic Variation on Proteasome Activity

**DOI:** 10.1101/2021.11.23.469794

**Authors:** Mahlon A. Collins, Randi R. Avery, Frank W. Albert

## Abstract

Protein degradation is an essential biological process that regulates protein abundance and removes misfolded and damaged proteins from cells. In eukaryotes, most protein degradation occurs through the stepwise actions of two functionally distinct entities, the ubiquitin system and the proteasome. Ubiquitin system enzymes attach ubiquitin to cellular proteins, targeting them for degradation. The proteasome then selectively binds and degrades ubiquitinated substrate proteins. Genetic variation in ubiquitin system genes creates heritable differences in the degradation of their substrates. However, the challenges of measuring the degradative activity of the proteasome independently of the ubiquitin system in large samples have limited our understanding of genetic influences on the proteasome. Here, using the yeast *Saccharomyces cerevisiae*, we built and characterized reporters that provide high-throughput, ubiquitin system-independent measurements of proteasome activity. Using single-cell measurements of proteasome activity from millions of genetically diverse yeast cells, we mapped 15 loci across the genome that influence proteasomal protein degradation. Twelve of these 15 loci exerted specific effects on the degradation of two distinct proteasome substrates, revealing a high degree of substrate-specificity in the genetics of proteasome activity. Using CRISPR-Cas9-based allelic engineering, we resolved a locus to a causal variant in the promoter of *RPT6*, a gene that encodes a subunit of the proteasome’s 19S regulatory particle. Our results reveal the complex genetic architecture of proteasome activity and suggest that genetic influences on the proteasome may be an important source of variation in the many cellular and organismal traits shaped by protein degradation.

**Author Summary:** Protein degradation controls the abundance of cellular proteins and serves an essential role in protein quality control by eliminating misfolded and damaged proteins. In eukaryotes, most protein degradation occurs in two steps. The ubiquitin system first targets proteins for degradation by attaching ubiquitin to them. The proteasome then selectively binds and degrades ubiquitinated proteins. Understanding how individual genetic differences affect the activity of the proteasome could improve our understanding of the many traits influenced by protein degradation. However, most assays that measure proteasomal protein degradation are not suitable for use in large samples or are affected by changes in the activity of the ubiquitin system. Using yeast, we built reporters that provide high-throughput measurements of proteasome activity independently of the ubiquitin system. We used measurements of proteasome activity from millions of live, single cells to identify regions of the genome with DNA variants that affect proteasomal protein degradation. We identified 15 such regions, showing that proteasome activity is a genetically complex trait. Using genome engineering, we found that one locus contained a variant in the promoter of a proteasome subunit gene that affected the activity of the proteasome towards multiple substrates. Our results demonstrate that individual genetic differences shape proteasome activity and suggest that these differences may contribute to variation in the many traits regulated by protein degradation.

## Introduction

Protein degradation helps maintain protein homeostasis by regulating protein abundance and eliminating misfolded and damaged proteins from cells. The primary protein degradation pathway in eukaryotes is the ubiquitin-proteasome system (UPS). The UPS consists of two functionally distinct components, the ubiquitin system and the proteasome^1–4^. Ubiquitin system enzymes bind degradation-promoting signal sequences (termed “degrons”^5^) in proteins, targeting bound substrate proteins for degradation by covalently attaching chains of the small protein ubiquitin (Figure 1A)^2, 3, 6, 7^. The proteasome then degrades polyubiquitinated proteins using two elements, the 19S regulatory particle and the 20S core particle^1, 8, 9^. The 19S regulatory particle selectively binds polyubiquitinated proteins^4, 10^ then deubiquitinates, unfolds, and translocates them to the 20S core particle, which degrades proteins to short peptides^11^(Figure 1A). The UPS is responsible for 70-80% of intracellular protein degradation^4, 12^ and influences the abundance of much of the proteome^13–15^. Therefore, UPS activity must be precisely and dynamically regulated at the levels of (1) substrate targeting by the ubiquitin system^16–18^ and (2) proteasomal protein degradation^19, 20^. Imbalances between UPS activity and the proteolytic needs of the cell adversely impact cellular viability and are associated with a diverse array of human diseases, including cancers, immune disorders, metabolic syndromes, and neurodegenerative diseases^3, 20–23^. Thus, determining the factors that create variation in substrate targeting by the ubiquitin system and proteasomal protein degradation could improve our understanding of the many traits influenced by protein degradation.

**Figure 1:**
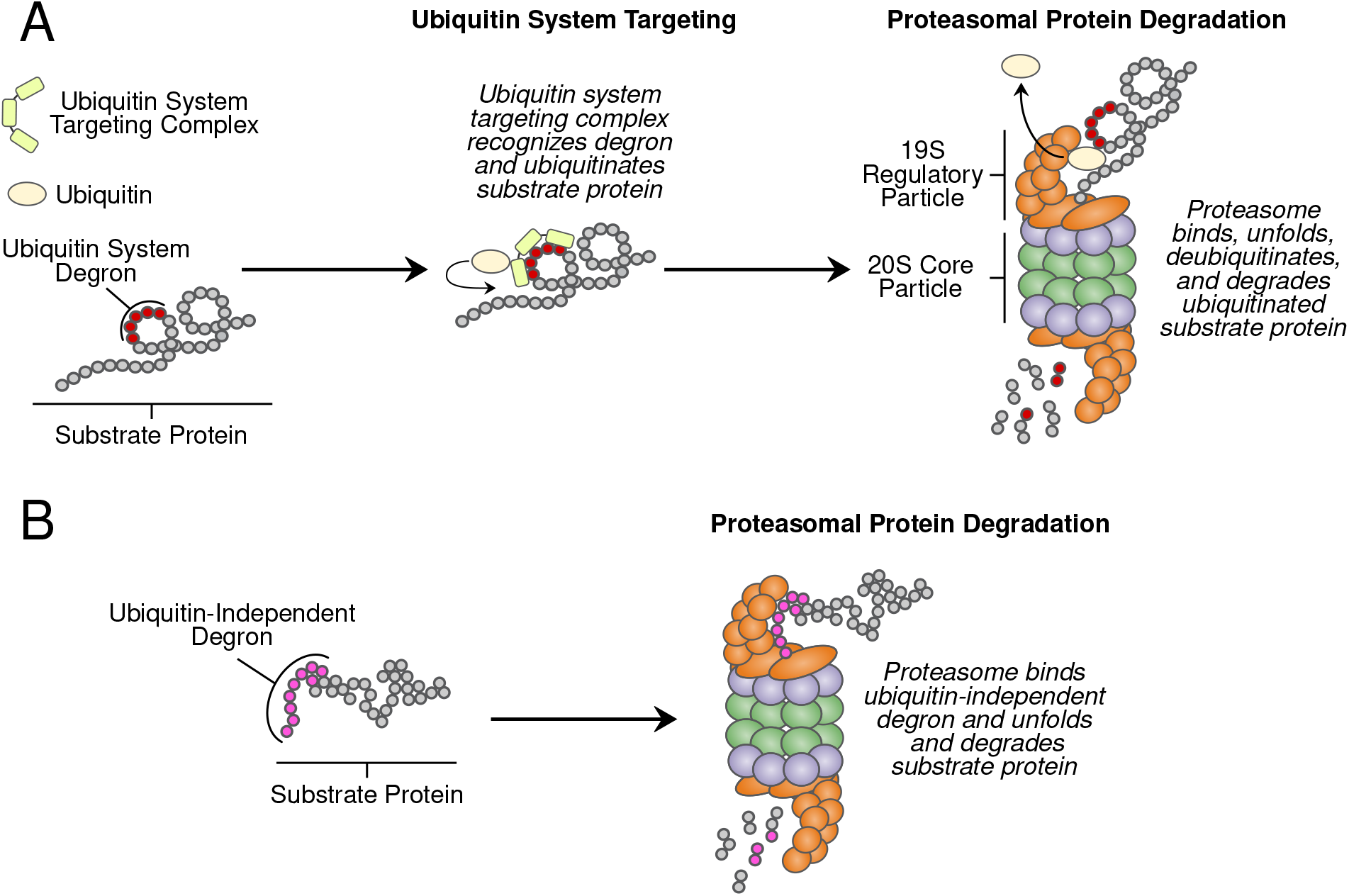
UPS protein degradation. A. UPS protein degradation resulting from (1) ubiquitin system targeting followed by (2) proteasomal protein degradation. B. Proteins with ubiquitin-independent degrons are directly bound and degraded by the proteasome without ubiquitin system targeting.

Until recently, it was largely unknown how individual genetic differences affect UPS protein degradation. To begin to address this question, we mapped genetic influences on the N-end Rule, a UPS pathway that recognizes degrons in protein N-termini (termed “N-degrons”^5, 24^). Our results showed that UPS activity is a genetically complex trait, shaped by variation throughout the genome^25^. Some of the largest genetic effects on N-end rule substrates resulted from variation in ubiquitin system genes. In particular, genes whose products process (*NTA1*) and recognize N-degrons (*UBR1* and *DOA10*) and ubiquitinate substrates (*UBC6*) each contained multiple causal variants that altered UPS activity, often in an N-degron-specific manner^25^. Thus, individual genetic differences in the ubiquitin system are an important source of substrate-specific variation in UPS protein degradation.

We do not know whether genetic effects on the proteasome are as prominent as those on the ubiquitin system. Our understanding of genetic influences on proteasome activity is largely limited to the clinical consequences of variation in proteasome genes. Missense mutations in several proteasome genes that alter proteasome activity cause a spectrum of heritable disease phenotypes, including intellectual disability^26^, lipodystrophy^27, 28^, cataracts^29^, recurrent fever^30^, and morphological abnormalities^31^. Variation in proteasome genes has also been linked to multiple common diseases, including myocardial infarction^32^, stroke^33^, type 2 diabetes^34, 35^, and cancer^36, 37^. However, these mutations and polymorphisms were identified through targeted sequencing of a subset of proteasome genes, leaving us with a biased, incomplete view of genetic influences on proteasome activity. Genome-wide association studies have linked variation in the vicinity of proteasome genes to a variety of organismal phenotypes^38–41^. However, these studies have neither fine-mapped the individual causal variants for these loci nor determined whether they alter proteasome activity.

A related question is whether variant effects on proteasome activity result in similar changes in the degradation of distinct proteasome substrates. Variation in protein half-lives spans several orders of magnitude^42–44^, in part as a result of proteasome-specific factors that are independent of the ubiquitin system, such as how readily proteins are bound, unfolded, and degraded by the proteasome. Substrate protein factors such as unstructured initiation region length^45–47^, biases in amino acid composition^48–50^, where in the protein degradation is initiated^45^, and the stability of a protein’s fold^48, 51^ can also alter how readily a specific protein is degraded by the proteasome. Moreover, the proteasome can exist in multiple configurations that can exhibit distinct preferences for individual protein substrates^52–56^. Thus, a systematic understanding of genetic effects on proteasome activity requires testing multiple proteasomal substrates with distinct sequence compositions.

Technical challenges have precluded a more systematic understanding of the genetics of proteasomal protein degradation. The effects of natural DNA polymorphisms are often subtle, necessitating large sample sizes for detection. Statistically powerful genetic mapping of cellular traits such as proteasome activity requires assays that can provide quantitative measurements from thousands of individuals^57^. At this scale, *in vitro* biochemical assays of proteasome activity are impractical. Several synthetic reporter systems can measure UPS activity *in vivo* with high throughput^58–60^. However, the output of these reporters reflects the activities of both the ubiquitin system and the proteasome. Thus, when using these systems to map genetic influences on UPS activity, variant effects on the ubiquitin system^25^ may mask or obscure specific effects on proteasomal protein degradation.

The proteasome degrades a handful of endogenous cellular proteins without ubiquitination, providing a means of directly measuring proteasome activity independently of the ubiquitin system (Figure 1B). These proteins contain ubiquitin-independent degrons, short peptides that promote rapid proteasomal degradation without ubiquitination^61–65^. Ubiquitin-independent degrons simultaneously function as proteasome recognition elements that engage the 19S regulatory particle and unstructured initiation regions for 20S core particle degradation (Figure 1B)^62, 64–69^. The degradation-promoting effect of these peptides is transferable; conjugating a ubiquitin-independent degron to a heterologous protein converts it to a short-lived, ubiquitin-independent proteasome substrate^64, 65, 67, 69, 70^. This property has been leveraged to create genetically encoded, high-throughput reporters of proteasome activity whose readout is independent of ubiquitin system activity^62, 70, 71^.

Here, we combined ubiquitin-independent degron-based proteasome activity reporters with our recently developed, statistically powerful mapping strategy to study the genetics of proteasome activity in the yeast *S. cerevisiae*. Our results reveal a polygenic genetic architecture of proteasome activity that is characterized by a high degree of substrate specificity. One locus contained a causal variant in the promoter of *RPT6*, a proteasome subunit gene, while other regions contained candidate causal genes with no known links to UPS protein degradation. Our results show that individual genetic differences are an important source of variation in proteasome activity that may contribute to the complex genetic basis of the many cellular and organismal traits influenced by protein degradation.

## Results

### Single-Cell Measurements Reveal Heritable Variation in Proteasome Activity

We sought to develop a reporter system capable of measuring proteasome activity independently of the ubiquitin system *in vivo* with high throughput and quantitative precision. To do so, we built a series of tandem fluorescent timers (TFTs), fusions of two fluorescent proteins with distinct spectral profiles and maturation kinetics^72, 73^. Our TFTs contained a faster-maturing green fluorescent protein (GFP^74^) and a slower-maturing red fluorescent protein (RFP^75^) (Figure 2A). The two fluorophores in the TFT mature at different rates and, as a result, the RFP / GFP ratio changes over time. If the TFT’s degradation rate is faster than the RFP’s maturation rate, the TFT’s output, expressed as the *−* log_2_ RFP / GFP ratio, is directly proportional to its degradation rate (Figure 2B). The TFT’s output is also independent of the TFT’s expression level^76^, making it possible to use TFTs in genetically diverse cell populations without confounding from genetic influences on reporter expression, which are expected in a genetically diverse cell population^14, 25, 76–79^.

**Figure 2:**
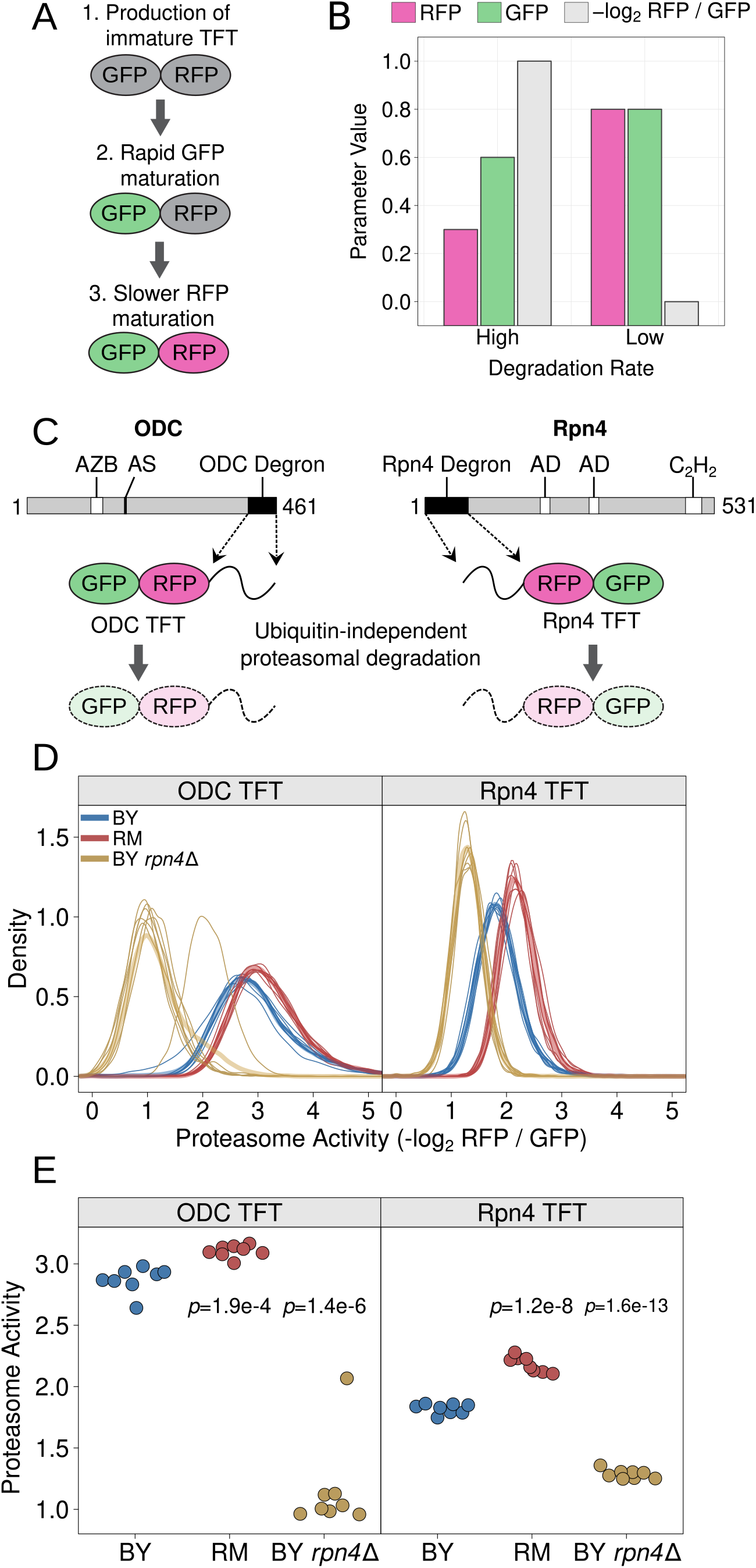
Design and characterization of proteasome activity reporters. A. Schematic of the production and maturation of a TFT. B. A bar plot created with simulated data shows how differences in a TFT’s degradation rate influence the reporter’s RFP and GFP levels, as well as the −log_2_ RFP / GFP ratio. C. Diagram of mouse ODC and yeast Rpn4 showing the location of each protein’s ubiquitin-independent degron. “AZB” = antizyme binding site, “AS” = active site, “AD” = transcriptional activation domain, “C_2_H_2_” = C_2_H_2_ zinc finger DNA binding domain. D. Density plots of proteasome activity from 10,000 cells for each of 8 independent biological replicates per strain per reporter for the indicated strains and TFTs. Thin, opaque lines show individual biological replicates and thicker, transparent lines show the group average for the indicated strains. E. The median from each biological replicate in D. is plotted as a stripchart. t-test p-values are shown for the indicated strain versus BY.

To relate the TFT’s output to proteasome activity, we fused the ubiquitin-independent degrons from the mouse ornithine decarboxylase (ODC) and yeast Rpn4 proteins to our TFTs (Figure 2C). When expressed in yeast, the mouse ODC degron is recognized, bound, and degraded by the proteasome^61, 67, 70^. This property has previously been used to measure proteasome activity *in vivo* in yeast cells^80^. We fused amino acids 410 through 461 of mouse ODC to the TFT’s C-terminus, consistent with the geometric requirements of the ODC degron^62^, to create the ODC TFT (Figure 2C). The Rpn4 protein contains a ubiquitin-independent degron in amino acids 1 to 80^64, 65^. We fused this sequence to the TFT’s N-terminus to create the Rpn4 TFT (Figure 2C). We reasoned that the distinct degron positions (C-and N-terminal), sequences, recognition mechanisms, and inferred 19S regulatory particle receptors^62, 64, 81^ would allow us to identify potential substrate-specific genetic effects on proteasome activity.

We characterized the ODC and Rpn4 TFTs in live, single cells by flow cytometry. We first evaluated the sensitivity of each TFT by comparing each TFT’s output in the BY laboratory strain and a BY strain lacking the *RPN4* gene (hereafter “BY *rpn4*Δ”). *RPN4* encodes a transcription factor for proteasome genes and deleting *RPN4* reduces proteasome activity^63, 68, 82^. Deleting *RPN4* strongly reduced the output from the ODC and Rpn4 TFTs in BY *rpn4*Δ (t-test *p* = 1.4e-6 and 1.6e-13, respectively; Figure 2D / E), showing that our TFTs provide sensitive *in vivo* measurements of proteasome activity. Consistent with previous reports^66, 69, 70^, in the BY strain the ODC TFT was more rapidly degraded than the Rpn4 TFT (t-test *p* = 6.9e-10, Figure 2D / E). Taken together, our results show that our TFTs provide quantitative, substrate-specific, *in vivo* readouts of proteasome activity.

To understand how natural genetic variation affects proteasome activity, we measured the output of the ODC and Rpn4 TFTs in two *Saccharomyces cerevisiae* strains. We compared BY, which is closely related to the S288C reference strain, and the genetically divergent vineyard strain, RM, whose genome differs from BY at an average at one out of every 200 base pairs^83^. The RM strain showed higher proteasome activity towards the ODC and Rpn4 TFTs than BY (t-test *p* = 1.9e-4 and 1.2e-8, respectively; Figure 2D / E). We observed a significant interaction between strain background and proteasome substrate such that the magnitude of the BY / RM strain difference was greater for the Rpn4 TFT than the ODC TFT (two-way ANOVA interaction *p* = 0.013). Together, these results show that individual genetic differences create heritable, substrate-specific variation in proteasome activity.

### Bulk Segregant Analysis Identifies Complex, Polygenic Influences on Proteasome Activity

To map genetic influences on proteasome activity, we used our ODC and Rpn4 TFTs to perform bulk segregant analysis, a statistically powerful genetic mapping method that compares large numbers of individuals with extreme values for a trait of interest selected from a genetically diverse population^25, 78, 79, 84, 85^. In our implementation, the method identifies quantitative trait loci (QTLs), regions of the genome with one or more DNA variants that influence proteasome activity. We created genetically diverse cell populations by mating BY strains harboring either the ODC or Rpn4 TFT with RM and sporulating the resulting diploids (Figure 3A). Using the resulting populations of haploid, genetically recombined progeny, we collected pools of 20,000 cells from the 2% tails of the proteasome activity distribution using fluorescence-activated cell sorting (FACS) (Figure 3B-E). We then whole-genome sequenced each pool to determine the allele frequency difference between the high and low UPS activity pools at each BY / RM DNA variant. At QTLs affecting proteasome activity, the allele frequencies will be significantly different between pools, while at unlinked loci the allele frequencies will be the same. We called significant QTLs using a logarithm of the odds (LOD) threshold previously determined to produce a 0.5% false discovery rate for TFT-based genetic mapping^25^ (see “Methods”) and retained only QTLs detected at genome-wide significance in both of two independent biological replicates. We determined the direction of QTL effects by computing the difference in RM allele frequency between the high and low proteasome activity pools at each QTL peak position. When this value is positive, the RM allele of the QTL results in higher proteasome activity, while negative values indicate QTLs where the RM allele decreases proteasome activity. We identified 11 QTLs for the ODC TFT and 7 QTLs for the Rpn4 TFT (Figure 4, Table 1). The distribution of proteasome activity QTL effect sizes, as reflected by the allele frequency difference between pools, was continuous and consisted predominantly of QTLs with small effects (Figure 4, Table 1). Together, our mapping results demonstrate that proteasome activity is a polygenic trait, shaped by variation throughout the genome.

**Figure 3:**
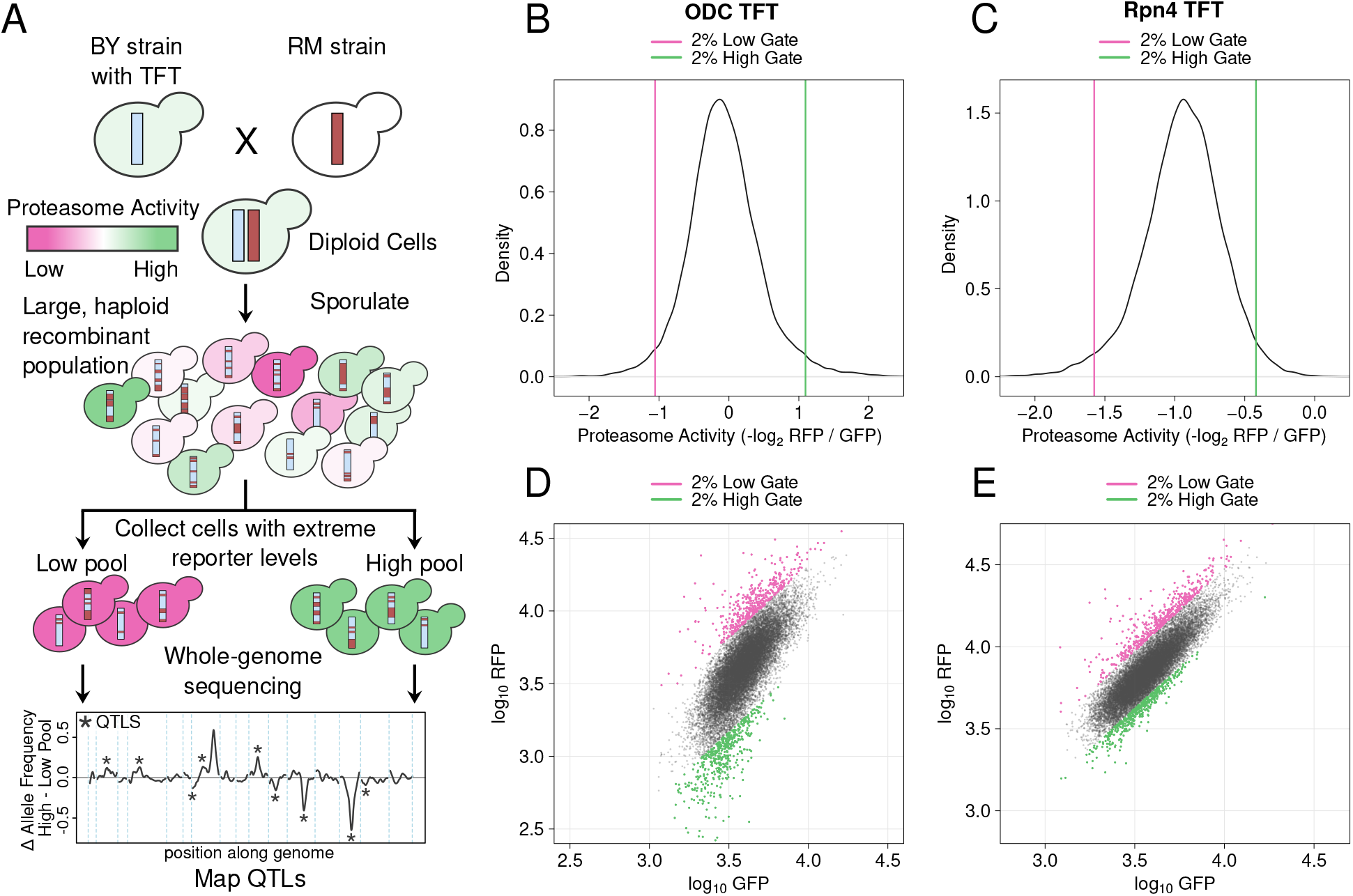
Mapping genetic influences on proteasome activity using bulk segregant analysis. A. Schematic of the experimental approach. B. / C. Proteasome activity distributions for the ODC TFT (B.) and Rpn4 TFT (C.). Vertical lines show the gates used to collect cells with extreme high or low proteasome activity. D. / E. Backplot of cells collected using the gates in B. / C. onto a scatter plot of GFP and RFP for the ODC (D.) and Rpn4 (E.) TFTs.

**Figure 4:**
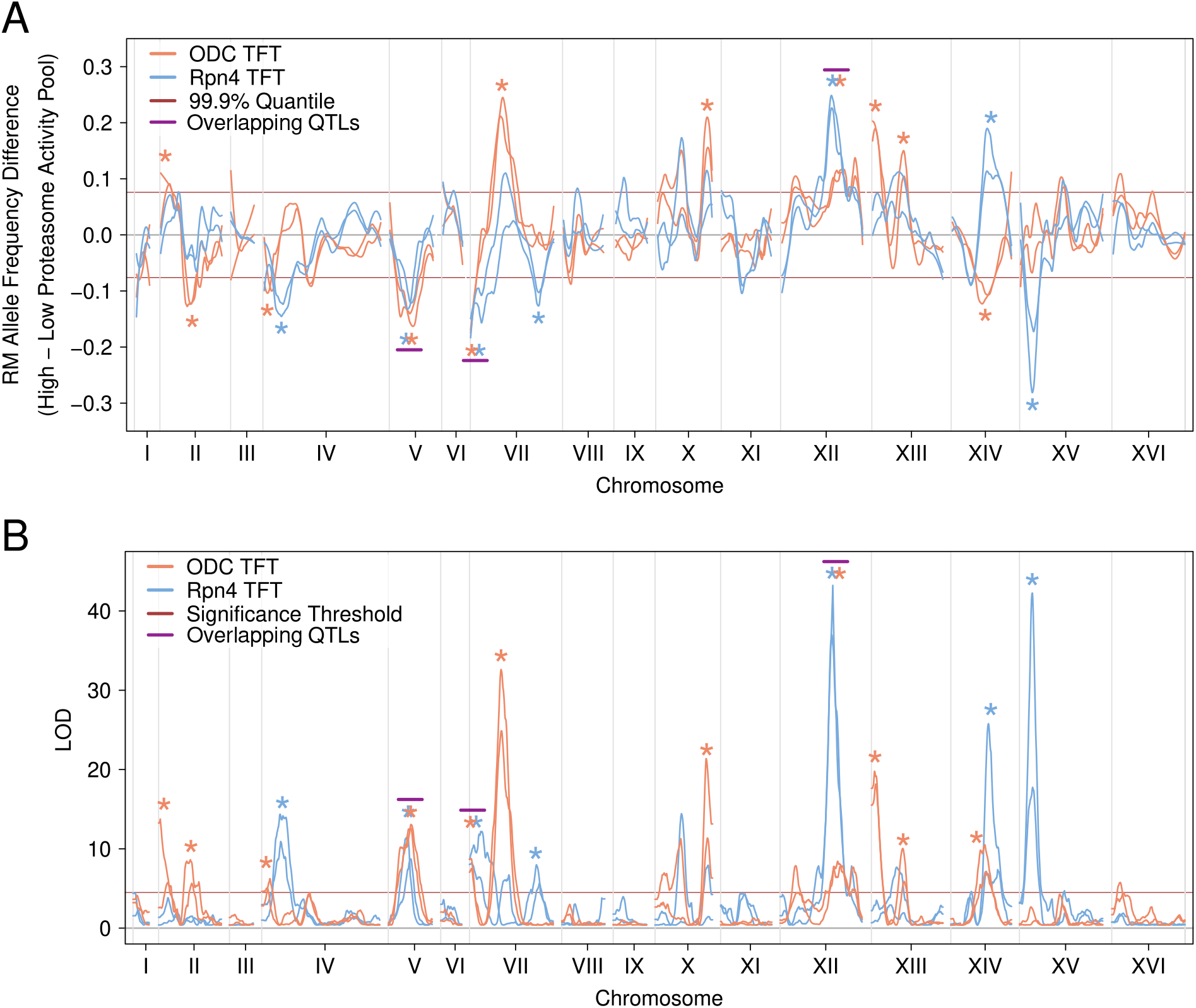
Proteasome activity QTLs detected with the ODC and Rpn4 TFTs. A. The line plot shows the loess-smoothed allele frequency difference between the high and low proteasome activity pools across the S. cerevisiae genome for each of two independent biological replicates per reporter. Asterisks denote QTLs, which are allele frequency differences exceeding an empirically-derived LOD score significance threshold (indicated in B.) in each of two independent biological replicates for a given reporter. The horizontal red lines denote an empirically-derived 99.9% quantile of the allele frequency difference. Magenta horizontal lines above pairs of asterisks denote QTLs detected with both TFTs with the same direction of effect, which are termed “overlapping QTLs”. B. As in A., but for the LOD score for proteasome activity QTLs. The red horizontal line denotes the LOD score significance threshold used to call QTLs at a 0.5% FDR.

**Table 1:**
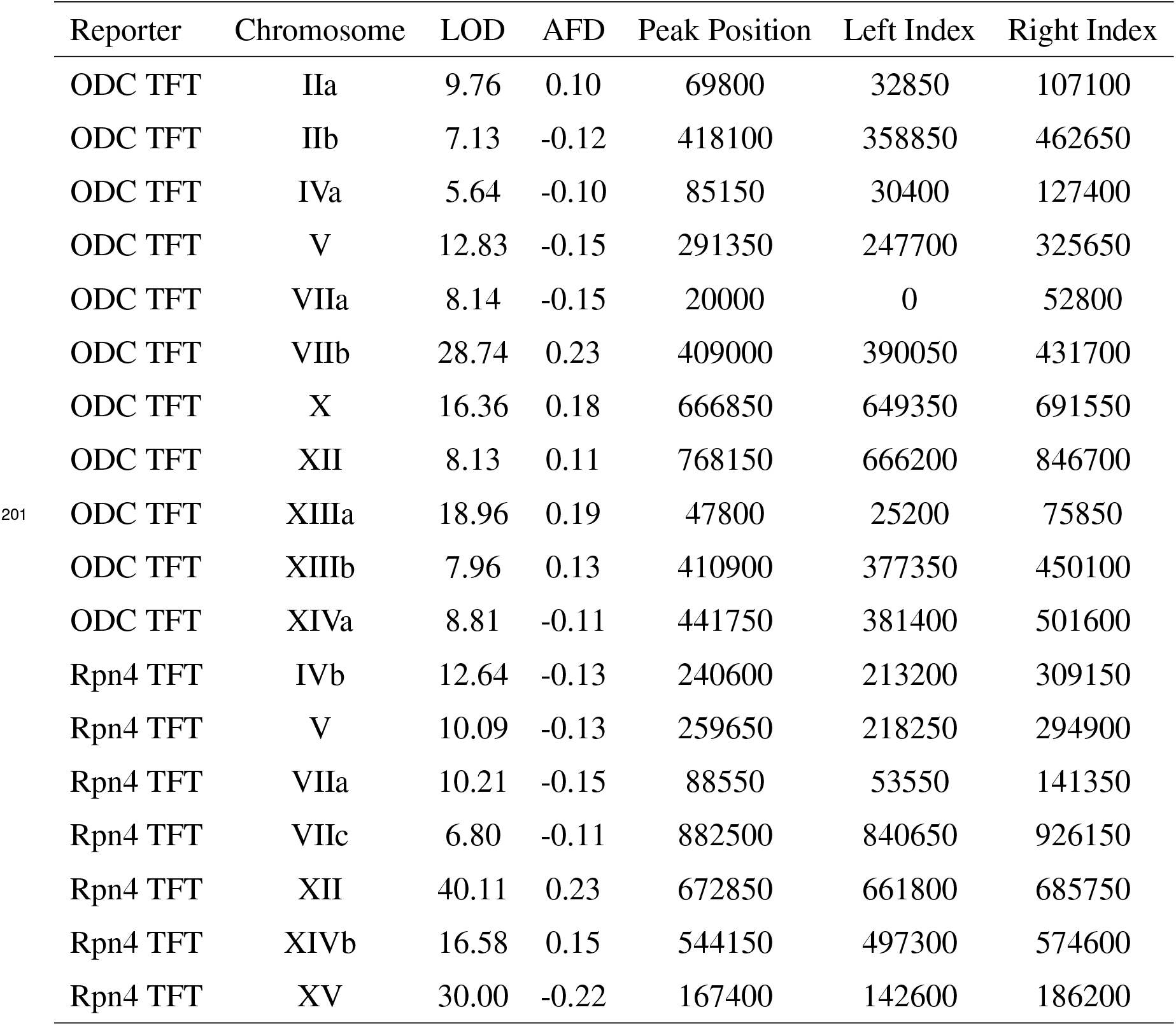
Proteasome activity QTLs detected with the ODC and Rpn4 TFTs. The table lists all detected QTLs, sorted first by reporter, then by chromosome. Lowercase letters following chromosome numbers are used to distinguish QTLs on the same chromosome. “LOD”, logarithm of the odds; “AFD”, RM allele frequency difference (high proteasome activity pool minus low proteasome activity pool) at the QTL peak position. “Peak Position”, “Left Index”, and “Right Index” refer to base pair positions on the indicated chromosome. Each number is the average value calculated from two independent biological replicates for a given QTL.

### Genetic Influences on Proteasome Activity are Predominantly Substrate-Specific

To study substrate specificity in the genetic architecture of proteasome activity, we evaluated the overlap in the sets of QTLs obtained with the ODC and Rpn4 TFTs. We defined overlapping QTLs as those whose peaks were within 100 kb of each other and had the same direction of effect. We then calculated the overlap fraction for the two sets of QTLs by dividing the number of overlapping QTLs by the number of overlapping QTLs plus the non-overlapping QTLs for each reporter. Only three proteasome activity QTLs, V, VIIA, and XII, overlapped between the sets of QTLs detected with the ODC and Rpn4 TFTs (overlap fraction = 0.2, Figure 4, Table 1), suggesting a high degree of substrate specificity.

To put this result in context, we examined overlap among our previously-described UPS N-end Rule activity QTLs^25^. The N-end Rule is divided into two primary branches based on how N-degrons are generated and recognized^86–89^. Ac/N-degrons are generated and recognized by a common set of molecular effectors^86^. Reflecting this, many QTLs for Ac/N-degrons affect all or a majority of the full set of Ac/N-degrons^25^. By contrast, Arg/N-degrons are created and recognized via molecular mechanisms that affect individual or small subsets of Arg/N-degrons^86^. Accordingly, QTLs for Arg/N-degrons tend to affect one or a minority of the set of Arg/N-degrons^25^. We computed the QTL overlap fraction among all pairs of Arg/N-degrons or Ac/N-degrons with at least 7 QTLs (to match the number of Rpn4 TFT QTLs detected here) using the criteria above. As expected, QTLs for Ac/N-degrons were detected with multiple reporters (median overlap fraction = 0.54; Figure 5A), while Arg/N-degron QTLs were more specific (median overlap fraction = 0.21; Figure 5A). The distributions of overlap fractions for Arg/N-degrons and Ac/N-degrons were highly distinct (Figure 5A), making them an ideal reference against which to gauge the substrate-specificity of proteasome activity QTLs.

**Figure 5:**
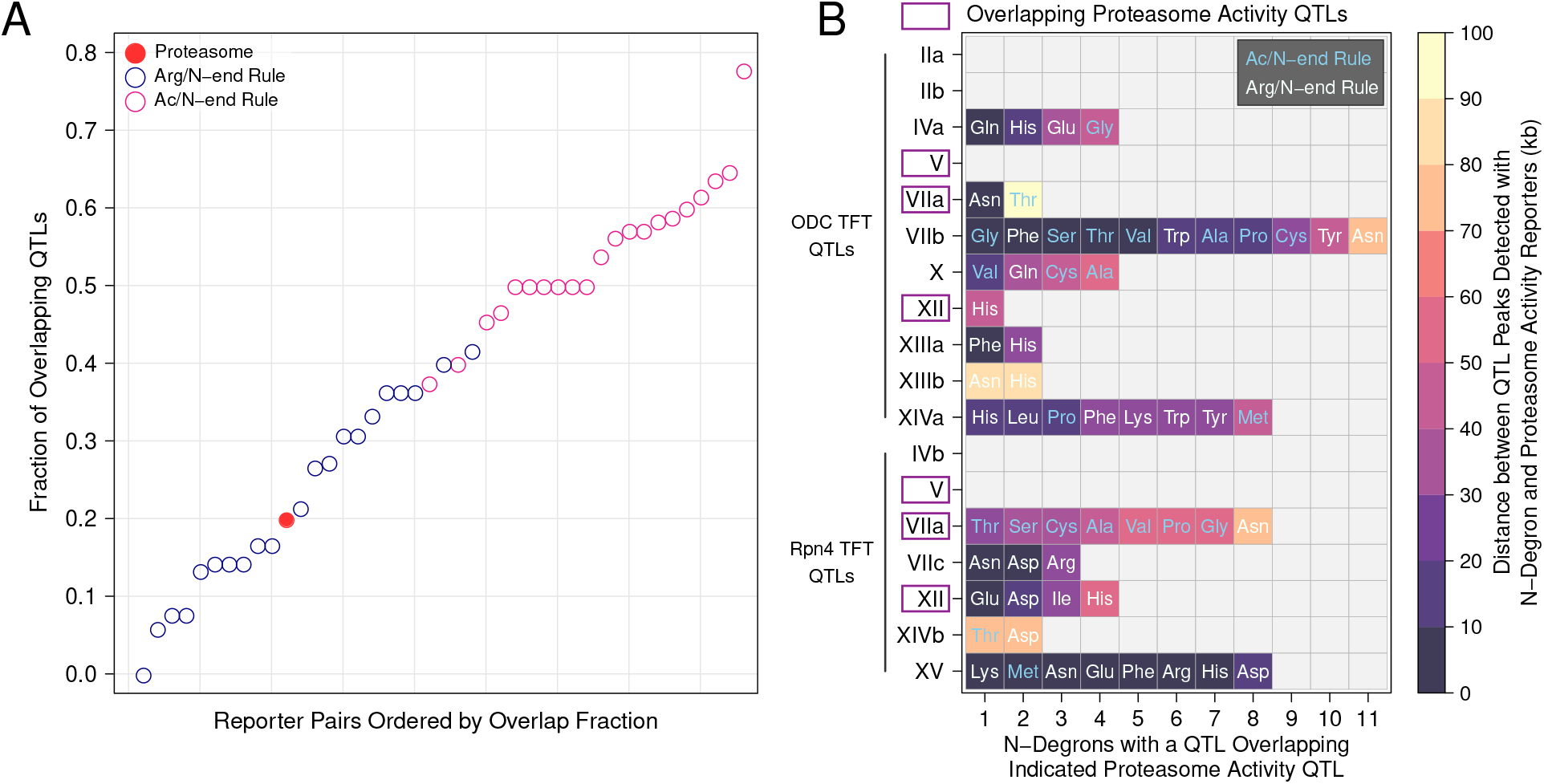
Overlap of N-end Rule and Proteasome Activity QTLs. A. Analysis of QTL overlap for proteasome activity, Arg/N-degron, and Ac/N-degron QTLs. For all pairs of reporters in the indicated reporter sets, we computed the overlap fraction as overlapping QTLs divided by total QTLs (overlapping QTLs plus reporter-specific QTLs). B. Overlap of proteasome activity and N-end Rule QTLs. The plot shows the number, identify, and N-end Rule branch of the N-degron QTLs that overlap proteasome activity QTLs on the y axis are ordered first by reporter then by chromosomal position and labeled as in Table 1. N-degrons on the x axis are ordered by the distance of their QTL’s peak position from the peak of the corresponding proteasome activity QTL detected with either the ODC or Rpn4 TFT.

The overlap fraction for the two sets of proteasome activity QTLs (0.2) was close to the median overlap for Arg/N-degrons (0.21, Figure 5A). Thus, genetic influences on proteasome activity are as substrate-specific as those on N-degrons that are engaged by a broad variety of molecular mechanisms in the ubiquitin system^86^. Overlap among the two sets of proteasome activity QTLs was considerably lower than that for the Ac/N-degrons (Figure 5A), which are generated and recognized via a common set of molecular effectors^86, 87^. Crucially, the current proteasome and previous N-end Rule QTLs were detected with a similar experimental design with similarly high statistical power. Therefore, these comparisons across datasets provide an estimate of substrate specificity that is immune to potential inflation from QTLs that truly affect multiple substrates but may appear to be substrate-specific because they happened to be detected with only one or a few reporters. The chromosome XIVa and XIVb QTLs, which occur at similar positions but have opposing effects on the degradation of the Rpn4 and ODC TFTs (Figure 4A), provide further evidence that genetic effects on proteasome activity are highly substrate-specific.

### Effects of Proteasome Activity QTLs on the UPS N-end Rule

We previously showed that four QTLs affecting the degradation of N-end Rule substrates contained causal variants in ubiquitin system genes^25^. As expected, these QTLs did not meet our criteria for overlap with any proteasome activity QTLs (Supplementary Table 1). However, many N-end Rule QTLs did not contain ubiquitin system genes, suggesting that they may result from genetic effects on processes unrelated to ubiquitin system targeting. To understand whether variation in N-end Rule activity could be explained by genetic effects on proteasome activity, we examined the overlap between the proteasome activity QTLs identified here and our previously-identified N-end Rule QTLs^25^. The set of N-end Rule QTLs comprises 149 QTLs detected with the 20 possible N-degron TFTs. However, many N-end Rule QTLs detected with distinct reporters overlap. To account for this, we applied our criteria for QTL overlap, which reduced the 149 N-end Rule QTLs detected with multiple reporters to 35 distinct, non-overlapping QTLs. Eleven proteasome activity QTLs overlapped one of these 35 N-end Rule QTLs (31%), suggesting that genetic effects on proteasome activity play a prominent role in shaping the activity of the UPS N-end Rule (Figure 5B).

Conversely, 4 of 15 proteasome activity QTLs did not overlap any N-end Rule QTLs, demonstrating that genetic variation can specifically alter the turnover of ubiquitin-independent proteasome substrates (Figure 5B). In particular, the chromosome V QTL altered the degradation of both the ODC and Rpn4 TFTs, but no N-end Rule TFTs, suggesting broad effects on ubiquitin-independent proteasomal degradation (Figure 5B). This agrees with previous findings that multiple factors specifically regulate the degradation of ubiquitin-independent proteasomal substrates, without affecting the degradation of ubiquitinated substrates^80^.

### Overlapping Proteasome Activity and N-end Rule QTLs Identify Candidate Causal Genes for Proteasome Activity

QTLs often span large intervals, complicating efforts to identify the underlying causal genes and variants. We reasoned that we could use overlapping proteasome activity and N-end Rule QTLs to more precisely estimate QTL peak positions and nominate candidate causal genes. To this end, we computed the overlaps between the sets of proteasome activity QTLs and N-end rule QTLs and used this information to identify candidate causal genes (Figure 5B). Two proteasome activity QTLs that were also detected with multiple N-degron TFTs occurred in genomic regions harboring variation that affects a multitude of traits in the BY / RM cross. The chromosome XIVa QTL was detected with the ODC TFT, 6 Arg/N-degron TFTs, and 2 Ac/N-degron TFTs (Figure 5B). The QTL’s average peak position at base pair 462,767 was located approximately 4.5 kb from the *MKT1* gene. *MKT1* encodes a multifunctional RNA binding protein involved in 3’ UTR-mediated RNA regulation^90, 91^. Variation at *MKT1* affects sporulation efficiency and growth^92, 93^. The *MKT1* locus also occurs in a gene expression QTL “hotspot” that influences the expression of thousands of genes^77, 78^ in the BY / RM cross. The chromosome XV QTL was detected with the Rpn4 TFT, 7 Arg/N-degron TFTs, and 1 Ac/N-degron TFT (Figure 5B). This set of QTL peaks clustered tightly at the average peak position of base pair 164,256. This position is approximately 7 kb away from *IRA2*, which encodes a negative regulator of RAS signaling^94^. Variation in *IRA2* affects the expression of thousands of genes in this cross of strains^95^ via multiple causal variants that interact epistatically^96^. The QTL intervals for the chromosome XIVa and XV QTLs do not contain any genes encoding proteasome subunits or proteasome assembly factors. Therefore, the QTLs at *MKT1* and *IRA2* illustrate that some genetic effects on proteasome activity likely result from complex, indirect molecular mechanisms involving altered gene expression.

The chromosome VIIb QTL detected with the ODC TFT had the highest number of overlapping N-end rule QTLs, with QTLs detected in the same region with 4 Arg/N-degron and 7 Ac/N-degron TFTs (Figure 5B). The high number of overlapping N-end Rule QTLs for both Arg/N-degrons and Ac/N-degrons suggested that this QTL contained variation that broadly affects UPS protein degradation. The average chromosome VIIb QTL peak position at base pair 411,250 is within the *RPT6* open reading frame. *RPT6* encodes a subunit of the proteasome’s 19S regulatory particle, suggesting that this QTL influences proteasome activity via direct effects on a proteasome subunit.

### Proteasome Activity is Shaped by a Causal Variant in the *RPT6* Promoter

We selected the chromosome VIIb QTL for further experimental dissection. There are no missense *RPT6* variants between BY and RM. However, a non-coding variant occurs at base pair 411,461 (Figure 6A) in an intergenic region between *RPT6* and the adjacent *ALG13*, which encodes an enzyme involved in oligosaccharide biosynthesis. We hypothesized that this intergenic variant (hereafter, “*RPT6* -175”) was the causal nucleotide for the chromosome VIIb QTL.

**Figure 6:**
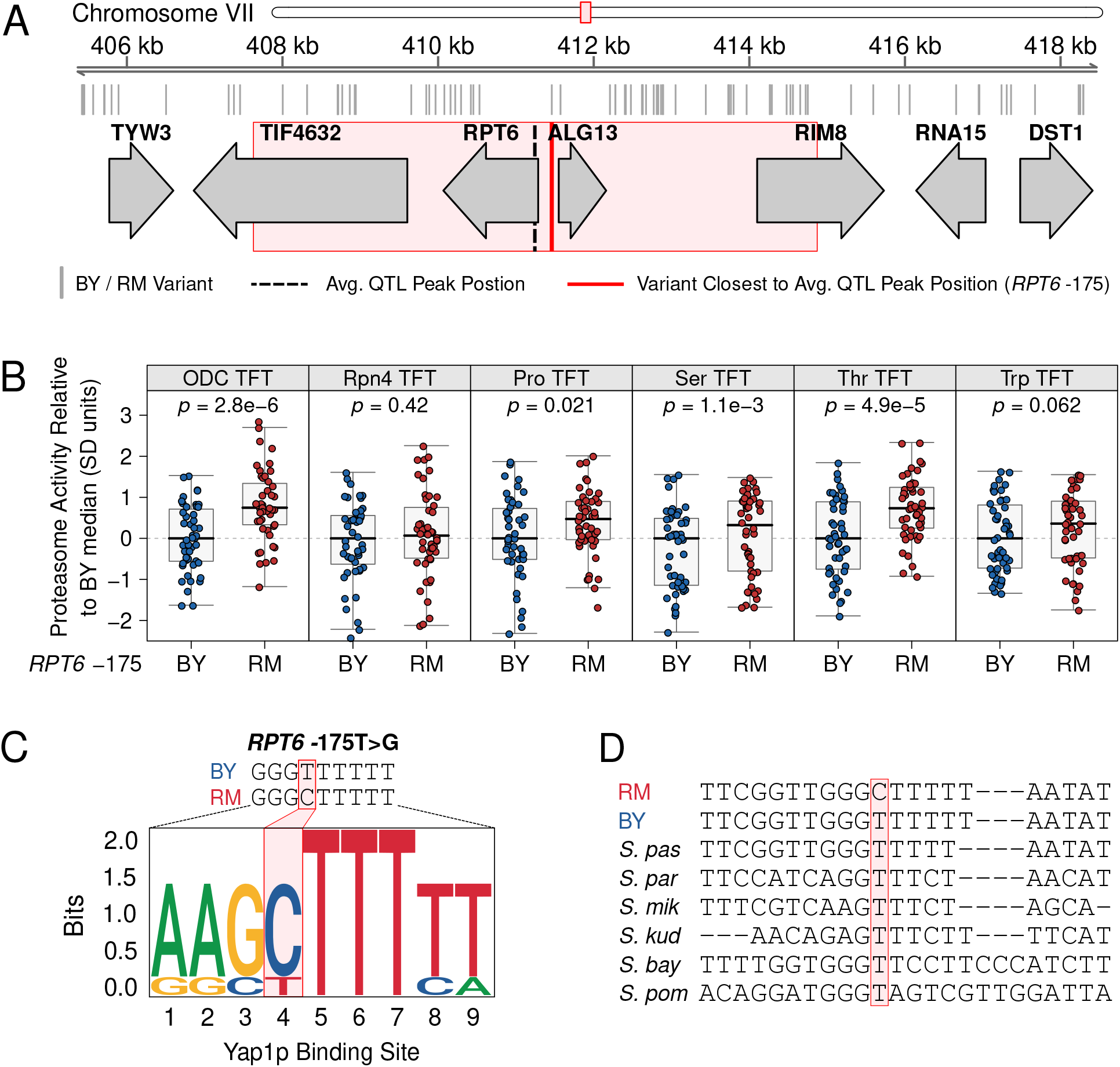
Fine-mapping a causal variant for the chromosome VIIb QTL. A. Genomic interval for the chromosome VIIb QTL. The red box depicts the 95% confidence interval of the chromosome VIIb QTL peak position, which was calculated using the chromosome VIIb QTL intervals from the ODC and N-end Rule TFTs with which the QTL was detected. B. CRISPR-Cas9 was used to engineer strains containing either the BY or RM allele at RPT6 -175 and the variant’s effect on proteasome activity was measured using the ODC and Rpn4 ubiquitin-independent degron TFTs, as well as the proline (Pro), serine (Ser), and threonine (Thr) Ac/N-end TFTs, and the tryptophan (Trp) Arg/N-degron TFT. C. Sequence logo for the predicted Yap1p binding site created by the RM allele of RPT6 -175. D. Multi-species alignment of the RPT6 promoter. The RPT6 -175 is highlighted. “S. pas” = Saccharomyces pastorianus, “S. par” = Saccharomyces paradoxus, “S. mik” *=* Saccharomyces mikatae, “S. kud” *=* Saccharomyces kudriavzevii, “S. bay” *=* Saccharomyces bayanus, “S. pombe” *=* Saccharomyces pombe

To test the effect of *RPT6* -175, we used CRISPR-Cas9 to create BY strains with either the BY or RM alleles at *RPT6* -175. We tested the effect of the *RPT6* -175 RM allele on the ODC and Rpn4 TFTs, as well as a subset of Ac/N-degron and Arg/N-degron reporters with which the chromosome VIIb QTL was also detected. The *RPT6* -175 RM allele significantly increased proteasome activity towards the ODC TFT as compared to the BY *RPT6* -175 allele (*p* = 2.8e-6, Figure 6B). Consistent with our QTL mapping results, the *RPT6* -175 RM allele did not increase proteasome activity towards the Rpn4 TFT (*p* = 0.42, Figure 6B). The *RPT6* -175 RM allele significantly increased the degradation of the proline, serine, and threonine Ac/N-degron TFTs, while its effect on the degradation of the tryptophan Arg/N-degron was not statistically significant (Figure 6B). These differences in how the *RPT6* -175 RM allele affects the Rpn4 and tryptophan TFTs compared to other reporters suggest that this allele exerts substrate-specific effects on proteasome activity. The Rpn4 degron, in particular, is recognized by distinct 19S regulatory particle receptors from the other substrates tested here^64^ and may, therefore, be unaffected by *RPT6* -175.

The *RPT6* -175 variant occurs in an intergenic region with putative promoters for *RPT6* and the divergently oriented *ALG13*. While we cannot formally exclude that the effect of the *RPT6* -175 occurs via *ALG13*, there are no known links between *ALG13* and proteasome activity. Moreover, previous genetic mapping in this cross of yeast strains identified a QTL in this same region affecting *RPT6* mRNA abundance with the same direction of effect as the proteasome activity QTL^77^. Increasing the expression of individual proteasome subunits is a well-established mechanism for increasing proteasome activity^19, 97, 98^. Based on these observations, we conclude that the effects of *RPT6* -175 on proteasome activity likely result from increased *RPT6* expression.

To understand potential molecular mechanisms of the *RPT6* -175 RM allele, we scanned the BY and RM *RPT6* promoters for transcription factor binding sites^99^. The RM, but not BY, allele contains a putative binding site for Yap1p (Figure 6C). Yap1p is a stress-associated transcription factor that indirectly increases proteasome activity during cellular stress, in part, by increasing expression of the proteasome gene transcription factor *RPN4*^100–102^. A multi-species alignment of the *RPT6* promoter, showed that the *RPT6* -175 BY allele is highly conserved among yeast species (Figure 6D). The BY allele is also present in the ancestral Taiwanese *S. cerevisiae* isolate, further indicating that the *RPT6* -175 RM allele is derived. We then examined *RPT6* -175 allelic status in a global panel of 1,011 *S. cerevisiae* isolates^103^ to better understand its population characteristics and evolutionary origin. Overall, the *RPT6* -175 RM allele has a 33.7% population frequency. However, among the “Wine / European” clade that contains RM, the *RPT6* -175 RM allele has a population frequency of 91.6% (Supplementary Figure 1). No other clades have a comparably high *RPT6* -175 RM allele frequency (Supplementary Figure 1). Yap1p increases proteasome activity in response to a variety of cellular stressors, including ethanol stress^104^. Thus, the *RPT6* -175 RM allele may have arisen in the “Wine / European” as an adaptation to the wine-making environment. Our results demonstrate that natural populations harbor derived alleles that increase proteasome activity and suggest that these alleles may have arisen through adaptation to local environmental conditions.

## Discussion

Much of the proteome undergoes regulated turnover via proteasomal protein degradation^13–15^. Proteasome activity is thus a critical determinant of the abundance of individual proteins and, by extension, the functional state of the cell. Physiological variation in proteasome activity enables adaptation to changing internal and external cellular environments, such as during cellular stress^97, 105, 106^, while pathological variation in proteasome activity is linked to a diverse array of human diseases^3, 20, 23, 107^. However, a full understanding of the factors that determine proteasome activity has remained elusive. In particular, the challenges of measuring proteasomal protein degradation in large samples has limited our understanding of the genetics of proteasome activity. By combining high-throughput proteasome activity reporters with a statistically powerful genetic mapping method, we have established individual genetic differences as an important source of variation in proteasome activity. Our results add to the emerging picture of the complex effects of genetic variation on protein degradation, which include widespread effects on the activity of the ubiquitin system^25^ and, as we show here, the proteasome.

This work provides several new insights into how individual genetic differences shape the activity of the proteasome. Previous studies identified rare mutations in proteasome genes as the cause of a variety of monogenic disorders^27–29, 31, 107, 108^. However, it was unclear to what extent these mutations are representative of genetic effects on proteasome activity. Our results suggest that disease-causing mutations and disease-linked polymorphisms with large effects on proteasome activity represent one extreme of a continuous distribution of variant effects on proteasome activity. Aberrant proteasome activity is a hallmark of numerous common human diseases^3, 20, 23^. Our results raise the possibility that the risk for these diseases may be subtly influenced by common variants that create heritable variation in proteasome activity. Our unbiased, genome-wide genetic mapping also identified QTLs containing no genes with previously-established connections to the regulation of proteasome activity. In particular, the chromosome XIVa and XV QTLs do not contain any genes encoding proteasome genes or proteasome assembly factors. Instead, the peaks of these QTLs center on *MKT1* and *IRA2*, which encode an RNA-binding protein and a RAS signaling regulator respectively, further highlighting the complexity of genetic effects on proteasome activity.

The proteasome activity QTLs we have identified add new insight into how genetic variation shapes the molecular effectors of cellular protein degradation. We recently mapped the genetics of the UPS N-end rule pathway and discovered multiple DNA variants that alter the activity of four functionally distinct components of the ubiquitin system^25^. Here, we extend this result by showing that genetic variation also shapes protein degradation through effects on the proteasome. Although many stimuli, such as protein misfolding or heat shock, cause coordinated changes in the activity of the ubiquitin system and the proteasome, recent work shows that these two systems can also be regulated independently and function autonomously of one another^19, 109^. For example, ubiquitination can initiate events besides proteasomal protein degradation, including lysosomal protein degradation, altered protein subcellular localization, and signaling cascade activation^109–111^. Likewise, a number of cellular proteins are bound and degraded by the proteasome without modification by the ubiquitin system^66^. Thus, predicting how genetic variation shapes the turnover of individual proteins will require consideration of genetic effects on both the ubiquitin system and the proteasome.

Genetic effects on proteasome activity were largely substrate-specific. Such a result would be expected if individual genetic differences primarily affected substrate selection by the proteasome’s 19S regulatory particle. Efficient degradation of the proteasome substrates tested here and in our previous study^25^ require the proteasome’s 19S regulatory particle^64, 112^, which contains multiple substrate receptors. The ODC and Rpn4 degrons are likely bound by distinct 19S receptors. Although the 19S receptors for the ODC degron are not known, the observation that the ODC degron competes with polyubiquitinated proteins for 19S binding has been used to infer that it is primarily bound by the canonical 19S ubiquitin receptors, Rpn1, Rpn10, and Rpn13^62, 81^. In contrast, the Rpn4 degron is bound by the 19S subunits Rpn2 and Rpn5^64^. Substrate selection is influenced by multiple factors, such as the subunit composition of the 19S regulatory particle and post-translational modification of 19S subunits^1, 8, 113^. Variant effects on these processes and other factors affecting 19S function may alter substrate selection, creating substrate-specific effects on proteasomal protein degradation. Four proteasome activity QTLs were not detected with any N-end reporters (Figure 5B), likely reflecting genetic mechanisms that specifically affect the degradation of substrates with ubiquitin-independent degrons. Collectively, our results suggest that genetic effects on protein degradation primarily affect subsets of UPS substrates, either via ubiquitin system targeting or at the substrate selection step preceding proteasomal protein degradation, rather than globally altering protein turnover.

Using CRISPR-Cas9 based allelic engineering, we resolved a QTL on chromosome VII to a causal nucleotide in the *RPT6* promoter. This region also harbors a QTL that influences *RPT6* expression^77^, suggesting the variant alters proteasome activity by altering *RPT6* expression. This mechanism is consistent with previous results showing that increasing the expression of individual proteasome subunits can increase proteasome activity. For example, increased expression of the 19S subunit *PSMD11* increases proteasome activity in human embryonic stem cells, which helps them maintain an undifferentiated stem cell identity^98^. In yeast, overexpression of the *PRE9* gene, which encodes a subunit of the 20S core particle, increases proteasome activity and promotes resistance to cellular stress^97^. Cells employ diverse mechanisms to monitor and degrade non-stoichoimetric subunits of protein complexes^114, 115^, raising the question of how increasing the expression of individual proteasome subunits increases proteasome activity. Rpt6 subunits not stably incorporated into proteasomes are protected from degradation by the chaperone proteasome associated assembly factor 1 (PAAF1)^115, 116^. Thus, PAAF1 association with Rpt6 creates a stable Rpt6 pool that can be used to rapidly drive proteasome assembly, leading to increased proteasome activity.

We have developed a generalizable strategy for mapping genetic effects on proteasomal protein degradation with high statistical power. The elements in our reporters function in many other eukaryotic organisms, including human cells^64, 67, 76^. Deploying the reporter systems developed here in genetically diverse cell populations may provide new insights into the genetic basis of a variety of cellular and organismal traits, including the many diseases marked by aberrant proteasome activity.

## Materials and Methods

### Tandem Fluorescent Timer (TFT) Reporters of Proteasome Activity

We used TFTs, fusions of two fluorescent proteins with distinct spectral profiles and maturation kinetics, to measure proteasome activity. The most common TFT implementation consists of a faster-maturing green fluorescent protein (GFP) and a slower-maturing red fluorescent protein (RFP)^72, 73, 76, 117^. Because the two fluorescent proteins mature at different rates, the RFP / GFP ratio changes over time. If the TFT’s degradation rate is faster than the RFP’s maturation rate, the negative log_2_ RFP / GFP ratio is directly proportional to the TFT’s degradation rate^72, 76^. The RFP / GFP ratio is also independent of the TFT’s expression level,^72, 76^, enabling high-throughput, quantitative measurements of TFT turnover in genetically diverse cell populations^25, 76^. All TFTs in the present study contained superfolder GFP (sfGFP)^74^ and the RFP mCherry^75^ separated by an unstructured 35 amino acid peptide sequence to minimize fluorescence resonance energy transfer^76^.

To measure proteasome activity with our TFTs, we fused the ubiquitin-independent degrons from the mouse ornithine decarboxylase (ODC) and yeast Rpn4 proteins to our sfGFP-mCherry TFTs. ODC, an enzyme involved in polyamine biosynthesis, contains a ubiquitin-independent degron in its C-terminal 37 amino acids^61, 62, 70, 118^. Rpn4, a transcription factor for proteasome genes, contains a ubiquitin-independent degron in its N-terminal 80 amino acids^63, 64, 68^. Both degrons are recognized and bound by the 19S regulatory particle without ubiquitin conjugation and function as unstructured initiation regions^46^ for 20S core particle degradation. Attaching either degron to a heterologous protein converts it into a short-lived proteasomal substrate with half-lives of approximately 5 minutes for the ODC degron and 20 minutes for the Rpn4 degron^66, 69, 70^. The ODC and Rpn4 degron sfGFP-mCherry TFTs thus provide direct, quantitative, substrate-specific readouts of proteasome activity.

We used a previously described approach^25^ to construct TFT reporters and yeast strains harboring TFTs. Each TFT contained the constitutively active *TDH3* promoter, the *ADH1* terminator, sfGFP, mCherry, and the KanMX selection module^119^. TFTs were constructed so that the ubiquitin-independent degron was immediately adjacent to mCherry (Figure 2C), consistent with established guidelines for optimizing TFT function^73^. We used BFA0190 as the plasmid backbone for all TFT plasmids. BFA0190 contains 734 bp of sequence upstream and 380 bp of sequence downstream of the *LYP1* ORF separated by a SwaI restriction site. We inserted TFT reporters into BFA0190 by digesting the plasmid with SwaI and inserting TFT components between the *LYP1* flanking sequences using isothermal assembly cloning (Hifi Assembly Cloning Kit; New England Biolabs [NEB], Ipswich, MA, USA). The 5’ and 3’ *LYP1* flanking sequences in each TFT plasmid contain natural SacI and BglII restriction sites, respectively. We produced linear DNA transformation fragments by digesting TFT-containing plasmids with SacI and BglII and gel purifying the fragments (Monarch Gel Purification, NEB). Genomic integration of each linear transformation fragment results in deletion of the *LYP1* gene, allowing selection for TFT integration at the *LYP1* locus using the toxic amino acid analogue thialysine (S-(2-aminoethyl)-L-cysteine hydrochloride)^120–122^ and G418^119^. All plasmids used in this study are listed in Supplementary Table 2.

### Yeast Strains and Handling

#### Yeast Strains

We used two genetically divergent *Saccharomyces cerevisiae* yeast strains for characterizing our proteasome activity TFTs and mapping genetic influences on proteasome activity. The haploid BY strain (genotype: *MATa his3*Δ *ho*Δ) is a laboratory strain that is closely related to the *S. cerevisiae* S288C reference strain. The haploid RM strain is a vineyard isolate with genotype *MATα can1*Δ::STE2pr-SpHIS5 *his3*Δ::NatMX *AMN1-BY ho*Δ::HphMX URA3-FY. BY and RM differ, on average, at 1 nucleotide per 200 base pairs, such that approximately 45,000 single nucleotide variants (SNVs) between the strains can serve as markers in a genetic mapping experiment^78, 79, 83, 84^. We also engineered a BY strain lacking the *RPN4* gene (hereafter “BY *rpn4*Δ”) to characterize the sensitivity and dynamic range of our TFT reporters. We replaced the *RPN4* gene with the NatMX cassette, which confers resistance to the antibiotic nourseothricin^119^. To do so, we transformed BY with a DNA fragment created by PCR amplifying the NatMX cassette from plasmid from Addgene plasmid #35121 (a gift from John McCusker) using primers with 40 bp of homology to the 5’ upstream and 3’ downstream sequences of *RPN4* using the transformation procedure described below. Strain genotypes are presented in Table 2. Supplementary Table 3 lists the full set of strains used in this study.

**Table 2:** Strain genotypes

| Short Name | Genotype | Antibiotic Resistance | Auxotrophies |
| --- | --- | --- | --- |
| BY | *MATa his3Δ hoΔ* |  | histidine |
| RM | *MATα can1Δ::STE2pr-SpHIS5*<br>*his3Δ::NatMX hoΔ::HphMX* | clonNAT, hygromycin | histidine |
| BY *rpn4Δ* | *MATa his3Δ hoΔ rpn4Δ::NatMX* | clonNAT | histidine |

The media formulations for all experiments are listed in Table 3. Synthetic complete media powders (SC -lys and SC -his -lys -ura) were obtained from Sunrise Science (Knoxville, TN, USA). We added the following reagents at the following concentrations to yeast media where indicated: G418, 200 mg / mL (Fisher Scientific, Pittsburgh, PA, USA); ClonNAT (nourseothricin sulfate, Fisher Scientific), 50 mg / L; thialysine (S-(2-aminoethyl)-L-cysteine hydrochloride; MilliporeSigma, St. Louis, MO, USA), 50 mg / L; canavanine (L-canavanine sulfate, Millipore-Sigma), 50 mg / L.

**Table 3:** Media Formulations

| Media Name | Abbreviation | Formulation |
| --- | --- | --- |
| Yeast-Peptone-Dextrose | YPD | 10 g / L yeast extract<br>20 g / L peptone<br>20 g / L dextrose |
| Synthetic Complete | SC | 6.7 g / L yeast nitrogen base<br>1.96 g / L amino acid mix -lys<br>20 g / L dextrose |
| Haploid Selection | SGA | 6.7 g / L yeast nitrogen base<br>1.74 g / L amino acid mix -his -lys -ura<br>20 g / L dextrose |
| Sporulation | SPO | 1 g / L yeast extract<br>10 g / L potassium acetate<br>0.5 g / L dextrose |

#### Yeast Transformations

We used the lithium acetate / single-stranded carrier DNA / polyethyline glycol (PEG) method for all yeast transformations^123^. In brief, yeast strains were inoculated into 5 mL of YPD liquid medium for overnight growth at 30 °C. The next day, we diluted 1 mL of each saturated culture into 50 mL of fresh YPD and grew cells for 4 hours. Cells were washed in sterile ultrapure water and then in transformation solution 1 (10 mM Tris HCl [pH 8.0], 1 mM EDTA [pH 8.0], and 0.1 M lithium acetate). After each wash, we pelleted the cells by centrifugation at 3,000 rpm for 2 minutes in a benchtop centrifuge and discarded supernatants. After washing, cells were suspended in 100 *µ*L of transformation solution 1 along with 50 *µ*g of salmon sperm carrier DNA and 300 ng of transforming DNA and incubated at 30 °C for 30 minutes with rolling. Subsequently, 700 *µ*L of transformation solution 2 (10 mM Tris HCl [pH 8.0], 1 mM EDTA [pH 8.0], and 0.1 M lithium acetate in 40% PEG) was added to each tube, followed by a 30 minute heat shock at 42 °C. Transformed cells were then washed in sterile, ultrapure water, followed by addition of 1 mL of liquid YPD medium to each tube. Cells were incubated in YPD for 90 minutes with rolling at 30 °C to allow for expression of antibiotic resistance cassettes. We then washed the cells with sterile, ultrapure water and plated 200 *µ*L of cells on solid SC -lys medium with G418 and thialysine, and, for strains with the NatMX cassette, clonNAT. We single-colony purified multiple independent colonies (biological replicates) from each transformation plate for further analysis as indicated in the text. Reporter integration at the targeted genomic locus was verified by colony PCR^124^ using the primers listed in Supplementary Table 4.

#### Yeast Mating and Segregant Populations

We used a modified synthetic genetic array (SGA) methodology^121, 122^ to create populations of genetically variable, recombinant cells (“segregants”) for genetic mapping. BY strains with either ODC or Rpn4 TFTs were mixed with the RM strain on solid YPD medium and grown overnight at 30 °C. We selected for diploid cells (successful BY / RM matings) by streaking mixed BY / RM cells onto solid YPD medium containing G418, which selects for the KanMX cassette in the TFT in the BY strain, and clonNAT, which selects for the NatMX cassette in the RM strain. Diploid cells were inoculated into 5 ml of liquid YPD and grown overnight at 30 °C. The next day, cultures were washed with sterile, ultrapure water, and resuspended in 5 mL of SPO liquid medium (Table 3). We induced sporulation by incubating cells in SPO medium at room temperature with rolling for 9 days. After confirming sporulation by brightfield microscopy, we pelleted 2 mL of cells, which were then washed with 1 mL of sterile, ultrapure water, and resuspended in 300 *µ*L of 1 M sorbitol containing 3 U of Zymolyase lytic enzyme (United States Biological, Salem, MA, USA) to degrade ascal walls. Asci were digested for 2 hours at 30 °C with rolling. Spores were then washed with 1 mL of 1 M sorbitol, vortexed for 1 minute at the highest intensity setting, and resuspended in sterile ultrapure water. We confirmed the release of cells from asci by brightfield microscopy and plated 300 *µ*l of cells onto solid SGA medium containing G418 and canavanine. This media formulation selects for haploid cells with (1) a TFT via G418, (2) the *MATa* mating type via the *Schizosaccharomyces pombe HIS5* gene under the control of the *STE2* promoter (which is only active in *MATa* cells), and (3) replacement of the *CAN1* gene with *S. pombe HIS5* via the toxic arginine analog canavanine^121, 122^. Haploid segregants were grown for 2 days at 30 °C and harvested by adding 10 mL of sterile, ultrapure water and scraping the cells from each plate. Each segregant population cell suspension was centrifuged at 3000 rpm for 10 minutes and resuspended in 1 mL of SGA medium. We added 450 *µ*L of 40% (v / v) sterile glycerol solution to 750 *µ*L to each segregant culture and stored this mixture in screw cap cryovials at *−*80 °C. We stored 2 independent sporulations each of the ODC and Rpn4 degron TFT-containing segregants (derived from our initial matings) as independent biological replicates.

### Flow Cytometry and Fluorescence-Activated Cell Sorting

#### Flow Cytometry

We characterized our proteasome activity TFTs using flow cytometry. For all flow cytometry experiments, we inoculated yeast strains into 400 *µ*L of liquid SC -lys medium with G418 for overnight growth in 2 mL 96 well plates at 30 °C with 1000 rpm mixing on a MixMate (Eppendorf, Hamburg, Germany). The next day, 4 *µ*L of each saturated culture was inoculated into a fresh 400 *µ*L of G418-containing SC -lys media and cells were grown for an additional 3 hours prior to flow cytometry. We performed all flow cytometry experiments on an LSR II flow cytometer (BD, Franklin Lakes, NJ, USA) equipped with a 20 mW 488 nm laser with 488 / 10 and 525 / 50 filters for measuring forward and side scatter and sfGFP fluorescence, respectively, as well as a 40 mW 561 nm laser and a 610 / 20 filter for measuring mCherry fluorescence. Table 4 lists the parameters and settings for all flow cytometry and fluorescence-activated cell sorting (FACS) experiments. All flow cytometry data was analyzed using R^125^ and the flowCore R package^126^. We filtered each flow cytometry dataset to exclude all events outside of 10% *±* the median forward scatter (a proxy for cell size). This gating approach captured the central peak of cells in the FSC histogram and removed cellular debris, aggregates of multiple cells, and restricted our analyses to cells of the same approximate size^25^.

**Table 4:** Flow cytometry and FACS settings.

| Parameter | Laser Line (nm) | Laser Setting (V) | Filter |
| --- | --- | --- | --- |
| forward scatter (FSC) | 488 | 500 | 488/10 |
| side scatter (SSC) | 488 | 275 | 488/10 |
| sfGFP | 488 | 500 | 525/50 |
| mCherry | 561 | 615 | 610/20 |

For flow cytometry experiments related to reporter characterization, we recorded 10,000 cells each from 8 independent biological replicates per strain for the ODC and Rpn4 degron TFTs. We extracted the median from each independent biological replicate and used these values for statistical analyses. The statistical significance of between strain differences for the ODC and Rpn4 degron TFTs was assessed using a two-tailed t-test without correction for multiple testing. We used an ANOVA with strain (BY or RM) and reporter (ODC or Rpn4 degron TFT) as fixed factors to assess the statistical significance of the interaction of genetic background with reporter.

For flow cytometry experiments related to fine-mapping the chromosome VIIb QTL, we used the following procedures. We recorded 10,000 cells each from 12 independent biological replicates per strain (BY *RPT6* -175 BY and BY *RPT6* -175 RM) per guide RNA per reporter (ODC and Rpn4 TFTs, as well as proline, serine, threonine, and tryptophan N-degron TFTs). We observed that, consistent with previous results^25^, the output of the TFTs changed over the course of each flow cytometry experiment. We used a previously-described approach in which the residuals of a regression of the TFT’s output on time were used to correct for this effect^25, 79^. We then Z-score normalized the sets of median values for each reporter, setting the mean equal to the median of the BY *RPT6* -175 BY allele strain. The effect of the *RPT6* -175 genotype was assessed using a linear mixed model implemented in the R packages ‘lme4’^127^ and ‘lmertest’^128^ using *RPT6* -175 genotype and guide RNA as fixed effects and plate as a random effect.

#### Fluorescence-Activated Cell Sorting (FACS)

We used FACS to collect pools of segregant cells for genetic mapping by bulk segregant analysis^78, 79^. We thawed and inoculated segregant populations into 5 mL of SGA medium containing G418 and canavanine for overnight growth at 30 °C with rolling. The following morning, we diluted 1 mL of cells from each segregant population into a fresh 4 mL of SGA medium containing G418 and canavanine. Diluted segregant cultures were grown for 4 hours prior to sorting on a FACSAria II cell sorter (BD). Plots of side scatter (SSC) height by SSC width and forward scatter (FSC) height by FSC width were used to remove doublets from each sample and cells were further filtered to contain cells within *±* 7.5% of the central FSC peak. We empirically determined that this filtering approach excluded cellular debris and aggregates while retaining the primary haploid cell population. We also defined a fluorescence-positive population by retaining only those TFT-containing cells with sfGFP fluorescence values higher than negative control BY and RM strains without TFTs. We collected pools of 20,000 cells each from the 2% high and low proteasome activity tails (Figure 2B / C) from two independent biological replicates for each TFT. Pools of 20,000 cells were collected into sterile 1.5 mL polypropylene tubes containing 1 mL of SGA medium that were grown overnight at 30 °C with rolling. After overnight growth, we mixed 750 *µ*L of cells with 450 *µ*L of 40% (v / v) glycerol and stored this mixture in 2 mL 96 well plates at *−*80 °C.

### Genomic DNA Isolation, Library Preparation, and Whole-Genome Sequencing

To isolate genomic DNA from sorted segregant pools, we first pelleted 800 *µ*L of each pool by centrifugation at 3,700 rpm for 10 minutes. Supernantants were discarded and cell pellets were resuspended in 800 *µ*L of a 1 M sorbitol solution containing 0.1 M EDTA, 14.3 mM *β*-mercaptoethanol, and 500 U of Zymolyase lytic enzyme (United States Biological) to digest cell walls. Zymolyase digestions were carried out by resuspending cell pellets with mixing at 1000 rpm for 2 minutes followed by incubation for 2 hours at 37 °C. After completing the digestion reaction, we pelleted and resuspended cells in 50 *µ*L of phosphate-buffered saline. We then used the Quick-DNA 96 Plus kit (Zymo Research, Irvine, CA, USA) to extract genomic DNA according to the manufacturer’s protocol, including an overnight protein digestion in a 20 mg / mL proteinase K solution at 55 °C prior to loading samples onto columns. DNA was eluted from sample preparation columns using 40 *µ*L of DNA elution buffer (10 mM Tris-HCl [pH 8.5], 0.1 mM EDTA). DNA concentrations for each sample were determined with the Qubit dsDNA BR assay kit (Thermo Fisher Scientific, Waltham, MA, USA) in a 96 well format using a Synergy H1 plate reader (BioTek Instruments, Winooski, VT, USA).

We used genomic DNA from our segregant pools to prepare a short-read library for whole-genome sequencing on the Illumina Next-Seq platform using a previously-described approach^25, 78, 79^. The Nextera DNA library kit (Illumina, San Diego, CA, USA) was used according to the manufacturer’s instructions with the following modifications. We fragmented and added sequencing adapters to genomic DNA by adding 5 ng of DNA to a master mix containing 4 *µ*L of Tagment DNA buffer, 1 *µ*L of sterile molecular biology grade water, and 5 *µ*L of Tagment DNA enzyme diluted 1:20 in Tagment DNA buffer and incubating this mixture on a SimpliAmp thermal cycler using the following parameters (Thermo Fisher Scientific): 55 °C temperature, 20 *µ*L reaction volume, 10 minute incubation. We PCR amplified libraries prior to sequencing by adding 10 *µ*L of the tagmentation reaction to a master mix containing 1 *µ*L of an Illumina i5 and i7 index primer pair mixture, 0.375 *µ*L of ExTaq polymerase (Takara), 5 *µ*L of ExTaq buffer, 4 *µ*L of a dNTP mixture, and 29.625 *µ*L of sterile molecular biology grade water. To multiplex samples for sequencing, we generated all 96 possible index oligo combinations using 8 i5 and 12 i7 index primers. Libraries were PCR amplified on a SimpliAmp thermal cycler using the following parameters: initial denaturation at 95 °C for 30 seconds, then 17 cycles of 95 °C for 10 seconds (denaturation), 62 °C for 30 seconds (annealing), and 72 °C for 3 minutes (extension). The DNA concentration of each reaction was quantified using the Qubit dsDNA BR assay kit (Thermo Fisher Scientific). We pooled equimolar amounts of each sample, ran this mixture on a 2% agarose gel, and extracted and purified DNA in the 400 bp to 600 bp region using the Monarch Gel Extraction Kit (NEB) according to the manufacturer’s instructions.

The pooled library was submitted to the University of Minnesota Genomics Center (UMGC) for quality control assessment and Illumina sequencing. UMGC staff performed three quality control (QC) assays prior to sequencing. The PicoGreen dsDNA quantification reagent (Thermo Fisher Scientific) was used to determine library concentration, with a concentration *≥* 1 ng/*µ*L required to pass. The Tapestation electrophoresis system (Agilent Technologies, Santa Clara, CA, USA) was used to determine library size, with libraries in the range of 200 to 700 bp passing. Finally, the KAPA DNA Library Quantification kit (Roche, Basel, Switzerland) was used to determine library functionality, with libraries requiring a concentration *≥* 2 nM to pass. The submitted library passed each QC assay. The library was sequenced on a Next-Seq 550 instrument in mid-output, 75 bp paired-end mode, generating 153,887,828 reads across all samples, with a median of 9,757,090 and a range of 5,994,921 to 14,753,319 reads per sample. The mean read quality for all samples was *>* 30. The median read coverage of the genome was 21, with a range of 16 to 25 across all samples. Data will be deposited into the NIH Sequence Read Archive following publication.

### QTL Mapping

We used a previously-described approach to identify QTLs from our whole-genome sequencing data^25, 78, 79^. We initially filtered our raw reads to retain only those with a mean base quality score greater than 30. Filtered reads were aligned to the *S. cerevisiae* reference genome (sacCer3) with the Burroughs-Wheeler alignment tool^129^. We used samtools^130^ to first remove unaligned reads, non-uniquely aligned reads, and PCR duplicates, and then to produce vcf files containing coverage and allelic read counts at each of 18,871 high-confidence, reliable SNPs^57, 84^, with BY alleles as reference and RM alleles as alternative alleles.

QTLs were called from allele counts using the MULTIPOOL algorithm^131^. MULTIPOOL estimates a logarithm of the odds (LOD) score by calculating a likelihood ratio from two models. In the noncausal model, the locus is not associated with the trait and the high and low proteasome activity pools have the same frequency of the BY and RM alleles. In the causal model, the locus is associated with the trait, such that the BY and RM allele frequencies differ between pools. QTLs were defined as loci with a LOD *≥* 4.5. In a previous study^25^, we empirically determined that this threshold produces a 0.5% false discovery rate (FDR) for TFT-based genetic mapping by bulk segregant analysis. We used the following MULTIPOOL settings: bp per centiMorgan = 2,200, bin size = 100 bp, effective pool size = 1,000. As in previous studies^78, 79^, we excluded variants with allele frequencies higher than 0.9 or lower than 0.1^25, 78, 79^. QTL confidence intervals were defined as a 2-LOD drop from the QTL peak (the QTL position with the highest LOD value). We computed the RM allele frequency difference (ΔAF) between the high and low proteasome activity pools at each allele to visualize QTLs. We also used ΔAF at each QTL peak to determine the magnitude and direction of the QTL’s effect. When the RM allele frequency difference at a QTL is positive, the RM allele of the QTL is associated with higher proteasome activity. Negative RM allele frequency differences indicate QTLs where the RM allele is associated with lower proteasome activity. Because allele frequencies are affected by random counting noise, we smoothed allele frequencies along the genome using loess regression prior to calculating ΔAF for each sample.

### QTL Fine-Mapping By Allelic Engineering

We used CRISPR-Cas9 to edit the *RPT6* -175 locus in the BY strain. Guide RNAs (gRNAs) targeting *RPT6* were obtained from the CRISPR track of the UCSC Genome Browser^132^. To control for potential off-target edits by CRISPR-Cas9, we used two unique guide RNAs to engineer each allelic edit. We selected two gRNAs in the *RPT6* open-reading frame (ORF) based on their proximity to the *RPT6* -175 variant (PAM sequences 226 and 194 bp from *RPT6* -175), their CRISPOR specificity scores^133^ (100 each, where 100 is the highest possible predicted specificity), and their predicted cleavage scores^134^ (66 and 56, where *>* 55 indicates high predicted cleavage efficiency). We inserted each gRNA into a plasmid that expresses Cas9 under the control of the constitutively active *TDH3* promoter as follows. We digested backbone plasmid BFA0224^25^ with the restriction enzymes HpaI and BsmBI (New England Biolabs) to remove the backbone vector’s existing gRNA. The cut vector was gel purified using the Monarch Gel Purification kit (New England Biolabs) according to the manufacturer’s instructions. We then performed isothermal assembly cloning using the HiFi Assembly Kit with the gel purified vector backbone and oligos encoding each gRNA (OFA1198 or OFA1199; Supplementary Table 4) to create plasmids BFA0242 and BFA0243 (Supplementary Table 2). Plasmids were miniprepped from DH5*α E. coli* cells using the Monarch Plasmid Miniprep kit. The sequence identities of BFA0242 and BFA0243 were confirmed by Sanger sequencing.

We created repair templates for co-transformation with BFA0242 and BFA0243 as follows. We first extracted genomic DNA from BY and RM using the “10 minute prep” protocol^135^. Genomic DNA from each strain was used as a template for PCR amplification of the *RPT6* promoter using oligos OFA1204 and OFA1207 (Supplementary Table 4). To prevent Cas9 cutting after editing of the *RPT6* -175 locus, we introduced two synonymous substitutions into the *RPT6* ORF by converting the serine codons GGA and TCA to AGT at base pairs 22-24 and 49-51. Synonmous substitutions were introduced using splicing overlap by extension PCR^136^ with primers OFA1208 and OFA1209. Full repair templates were then amplified using either the BY or RM *UBR1* promoter and the BY *RPT6* ORF as templates in a splicing overlap extension by PCR reaction with primers OFA1204 and OFA1205 (Supplementary Table 4). The sequence identify of all repair templates was verified by Sanger sequencing.

To create BY strains with edited *RPT6* alleles, we co-transformed 150 ng of either plasmid BFA0242 or BFA0243 with 1.5 *µ*g of repair template using the transformation protocol above. The transformation reaction was streaked onto solid SC medium lacking histidine to select for the *HIS3* selectable marker in BFA0242 or BFA0243. Colonies from transformation plates were single-colony purified on solid medium lacking histidine, then patched onto solid YPD medium. To verify allelic edits, we performed colony PCR using oligos 1204 and 1206 (Supplementary Table 4). Reaction products were gel purified using the Monarch Gel Purification kit (New England Biolabs) and Sanger sequenced using oligos OFA1204 and OFA1206 to confirm both the sequence of the *RPT6* promoter and the synonymous substitutions in the *RPT6* ORF. Strains with the desired edits were then transformed to contain TFT reporters as indicated above. We tested 12 independent biological replicates per strain per guide RNA per TFT. For subsequent statistical analyses, we pooled strains with the same allelic edit engineered with unique guide RNAs.

### Data and Statistical Analysis

All data and statistical analyses were performed using R^125^. In all boxplots, the center line shows the median, the box bounds the first and third quartiles, and the whiskers extend to 1.5 times the interquartile range. DNA binding motifs in the *RPT6* promoter were assessed using the Yeast Transcription Factor Specificity Compendium database^99^. We inferred the allelic status of *RPT6* -175 by comparing the BY and RM alleles to a likely-ancestral Taiwanese strain. The frequency of the RM allele at *RPT6* -175 was calculated across and within clades of a global panel of 1,011 *S. cerevisiae* isolates^103^. Final figures and illustrations were made using Inkscape (version 0.92; Inkscape Project).

### Data and Materials Availability

Computational scripts used to process data, for statistical analysis, and to generate plots are available at: http://www.github.com/mac230/proteasome_QTL_paper Whole-genome sequencing data is in the process of being deposited into the NIH Sequence Read Archive. Yeast strains and plasmids used in this study are available on request. Correspondence should be addressed to FWA.

## Author Contributions

Conceptualization: MAC, FWA

Formal Analysis: MAC

Funding Acquisition: MAC, FWA

Investigation: MAC, RRA

Methodology: MAC, FWA

Resources: FWA

Supervision: MAC, FWA

Validation: MAC, RRA

Visualization: MAC

Writing - Original Draft: MAC

Writing - Review and Editing: MAC, FWA

## Acknowledgements

We thank the members of the Albert laboratory for feedback on the project and manuscript. We thank the University of Minnesota’s Flow Cytometry Resource and Genomics Center for their contributions to the project.

## Supporting Information

### Supplementary Figure

**Supplementary Figure 1.**
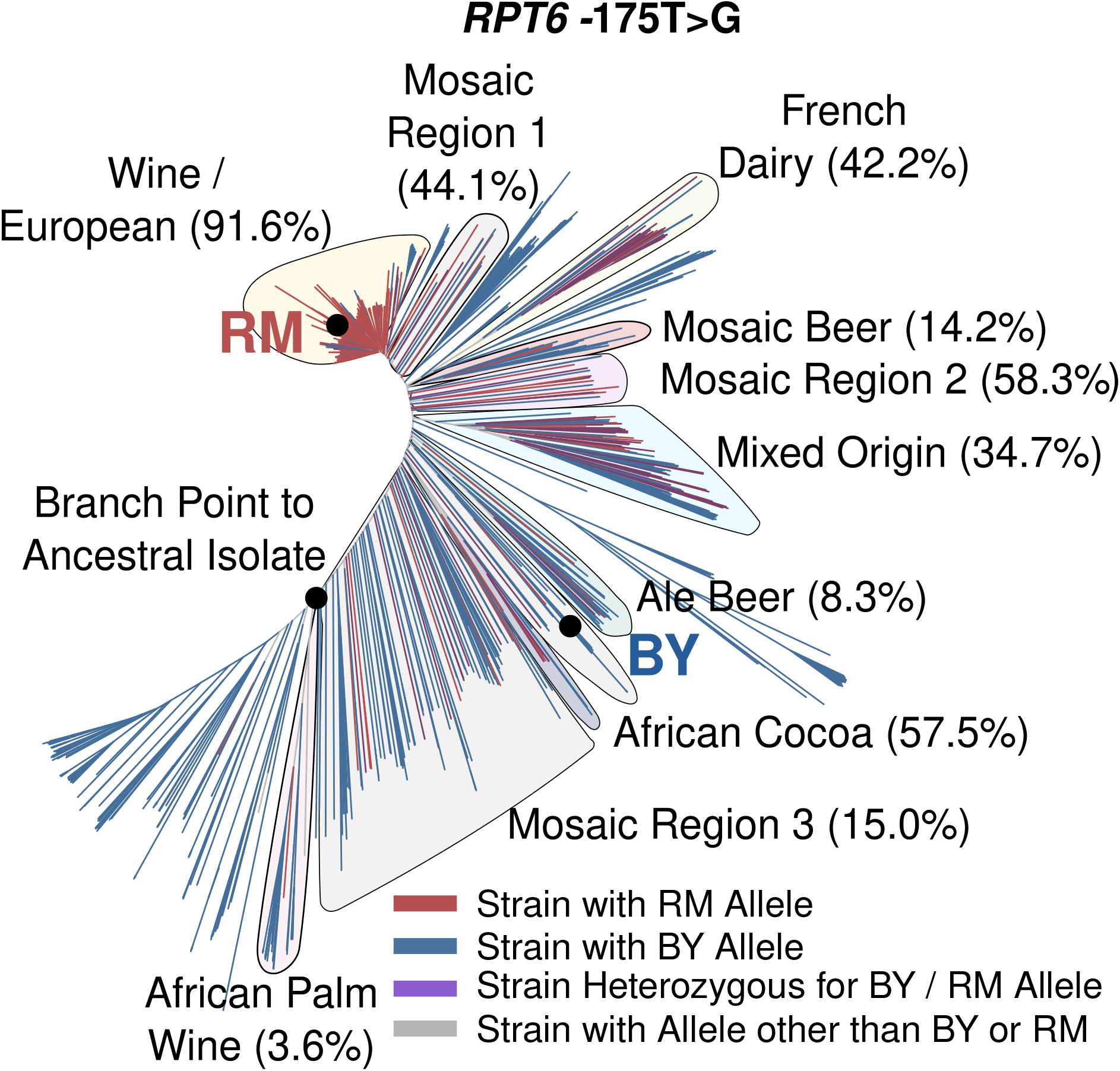
Tree diagram showing the distribution of the RPT6 -175 allele among a panel of 1,011 S. cerevisiae strains. Clades with the RPT6 -175 RM allele are indicated along with its frequency in that clade in parentheses.

### Supplementary Table Captions

**Supplementary Table 1.** *Overlap of proteasome activity QTLs with known causal genes for N-end Rule QTLs*.

**Supplementary Table 2.** *Plasmids used in the study*.

**Supplementary Table 3.** *Yeast strains used in the study*.

**Supplementary Table 4.** *Oligonucleotides used in the study*.

## References

1 D. Finley, H. D. Ulrich, T. Sommer, and P. Kaiser. The ubiquitin-proteasome system of Saccharomyces cerevisiae. Genetics, 192(2):319–360, Oct 2012.

2 A. Hershko and A. Ciechanover. The ubiquitin system. Annu Rev Biochem 67:425–479, 1998.

3 A. L. Schwartz and A. Ciechanover. The ubiquitin-proteasome pathway and pathogenesis of human diseases. Annu Rev Med, 50:57–74, 1999.

4 G. A. Collins and A. L. Goldberg. The Logic of the 26S Proteasome. Cell, 169(5):792–806, May 2017.

5 A. Varshavsky. Naming a targeting signal. Cell, 64(1):13–15, Jan 1991.

6 A. Ciechanover, A. Orian, and A. L. Schwartz. Ubiquitin-mediated proteolysis: biological regulation via destruction. Bioessays, 22(5):442–451, May 2000.

7 N. S. Abell, M. K. DeGorter, M. J. Gloudemans, E. Greenwald, K. S. Smith, Z. He, and S. B. Montgomery. Multiple causal variants underlie genetic associations in humans. Science, 375(6586):1247–1254, 03 2022.

8 J. Hanna and D. Finley. A proteasome for all occasions. FEBS Lett, 581(15):2854–2861, Jun 2007.

9 T. Inobe and A. Matouschek. Paradigms of protein degradation by the proteasome. Curr Opin Struct Biol, 24:156–164, Feb 2014.

10 O. Coux, K. Tanaka, and A. L. Goldberg. Structure and functions of the 20S and 26S proteasomes. Annu Rev Biochem, 65:801–847, 1996.

11 A. F. Kisselev, T. N. Akopian, K. M. Woo, and A. L. Goldberg. The sizes of peptides generated from protein by mammalian 26 and 20 S proteasomes. Implications for understanding the degradative mechanism and antigen presentation. J Biol Chem, 274(6):3363–3371, Feb 1999.

12 J. Zhao, B. Zhai, S. P. Gygi, and A. L. Goldberg. mTOR inhibition activates overall protein degradation by the ubiquitin proteasome system as well as by autophagy. Proc Natl Acad Sci U S A, 112(52):15790–15797, Dec 2015.

13 R. Christiano, H. Arlt, S. Kabatnik, N. Mejhert, Z. W. Lai, R. V. Farese, and T. C. Walther. A Systematic Protein Turnover Map for Decoding Protein Degradation. Cell Rep, 33(6):108378, 11 2020.

14 K. E. Kong, B. Fischer, M. Meurer, I. Kats, Z. Li, F. Rühle, J. D. Barry, D. Kirrmaier, V. Chevyreva, B. J. San Luis, M. Costanzo, W. Huber, B. J. Andrews, C. Boone, M. Knop, and A. Khmelinskii. Timer-based proteomic profiling of the ubiquitin-proteasome system reveals a substrate receptor of the GID ubiquitin ligase. Mol Cell, 81(11):2460–2476, 06 2021.

15 V. Solomon, S. H. Lecker, A. L. Goldberg, and A. L. Goldberg. The N-end rule pathway catalyzes a major fraction of the protein degradation in skeletal muscle. J Biol Chem, 273(39):25216–25222, Sep 1998.

16 S. E. Smith, M. Koegl, and S. Jentsch. Role of the ubiquitin/proteasome system in regulated protein degradation in Saccharomyces cerevisiae. Biol Chem, 377(7-8):437–446, 1996.

17 D. Kornitzer and A. Ciechanover. Modes of regulation of ubiquitin-mediated protein degradation. J Cell Physiol, 182(1):1–11, Jan 2000.

18 J. S. Bett. Proteostasis regulation by the ubiquitin system. Essays Biochem, 60(2):143–151, 10 2016.

19 R. S. Marshall and R. D. Vierstra. Dynamic Regulation of the 26S Proteasome: From Synthesis to Degradation. Front Mol Biosci, 6:40, 2019.

20 M. Schmidt and D. Finley. Regulation of proteasome activity in health and disease. Biochim Biophys Acta, 1843(1):13–25, Jan 2014.

21 R. Shringarpure and K. J. Davies. Protein turnover by the proteasome in aging and disease. Free Radic Biol Med, 32(11):1084–1089, Jun 2002.

22 C. Zheng, T. Geetha, and J. R. Babu. Failure of ubiquitin proteasome system: risk for neurodegenerative diseases. Neurodegener Dis, 14(4):161–175, 2014.

23 N. P. Dantuma and L. C. Bott. The ubiquitin-proteasome system in neurodegenerative diseases: precipitating factor, yet part of the solution. Front Mol Neurosci, 7:70, 2014.

24 A. Varshavsky. N-degron and C-degron pathways of protein degradation. Proc Natl Acad Sci U S A, 116(2):358–366, 01 2019.

25 Mahlon A. Collins, Gemechu Mekonnen, and Frank W. Albert. Variation in ubiquitin system genes creates substrate-specific effects on proteasomal protein degradation. bioRxiv, 2021.

26 J. de Ligt, M. H. Willemsen, B. W. van Bon, T. Kleefstra, H. G. Yntema, T. Kroes, A. T. Vultovan Silfhout, D. A. Koolen, P. de Vries, C. Gilissen, M. del Rosario, A. Hoischen, H. Scheffer, B. B. de Vries, H. G. Brunner, J. A. Veltman, and L. E. Vissers. Diagnostic exome sequencing in persons with severe intellectual disability. N Engl J Med, 367(20):1921–1929, Nov 2012.

27 A. K. Agarwal, C. Xing, G. N. DeMartino, D. Mizrachi, M. D. Hernandez, A. B. Sousa, L. Martínez de Villarreal, H. G. dos Santos, and A. Garg. PSMB8 encoding the B5i proteasome subunit is mutated in joint contractures, muscle atrophy, microcytic anemia, and panniculitis-induced lipodystrophy syndrome. Am J Hum Genet, 87(6):866–872, Dec 2010.

28 Y. Liu, Y. Ramot, A. Torrelo, A. S. Paller, N. Si, S. Babay, P. W. Kim, A. Sheikh, C. C. Lee, Y. Chen, A. Vera, X. Zhang, R. Goldbach-Mansky, and A. Zlotogorski. Mutations in proteasome subunit B type 8 cause chronic atypical neutrophilic dermatosis with lipodystrophy and elevated temperature with evidence of genetic and phenotypic heterogeneity. Arthritis Rheum, 64(3):895–907, Mar 2012.

29 A. Kröll-Hermi, F. Ebstein, C. Stoetzel, V. Geoffroy, E. Schaefer, S. Scheidecker, S. Bär, M. Takamiya, K. Kawakami, B. A. Zieba, F. Studer, V. Pelletier, C. Eyermann, C. Speeg-Schatz, V. Laugel, D. Lipsker, F. Sandron, S. McGinn, A. Boland, J. F. Deleuze, L. Kuhn, J. Chicher, P. Hammann, S. Friant, C. Etard, E. Krüger, J. Muller, U. Strähle, and H. Dollfus. Proteasome subunit PSMC3 variants cause neurosensory syndrome combining deafness and cataract due to proteotoxic stress. EMBO Mol Med, 12(7):e11861, 07 2020.

30 A. Brehm, Y. Liu, A. Sheikh, B. Marrero, E. Omoyinmi, Q. Zhou, G. Montealegre, A. Biancotto, A. Reinhardt, A. Almeida de Jesus, M. Pelletier, W. L. Tsai, E. F. Remmers, L. Kardava, S. Hill, H. Kim, H. J. Lachmann, A. Megarbane, J. J. Chae, J. Brady, R. D. Castillo, D. Brown, A. V. Casano, L. Gao, D. Chapelle, Y. Huang, D. Stone, Y. Chen, F. Sotzny, C. C. Lee, D. L. Kastner, A. Torrelo, A. Zlotogorski, S. Moir, M. Gadina, P. McCoy, R. Wesley, K. I. Rother, K. Rother, P. W. Hildebrand, P. Brogan, E. Krüger, I. Aksentijevich, and R. Goldbach-Mansky. Additive loss-of-function proteasome subunit mutations in CANDLE/PRAAS patients promote type I IFN production. J Clin Invest, 125(11):4196–4211, Nov 2015.

31 U. Tomaru, S. Takahashi, A. Ishizu, Y. Miyatake, A. Gohda, S. Suzuki, A. Ono, J. Ohara, T. Baba, S. Murata, K. Tanaka, and M. Kasahara. Decreased proteasomal activity causes age-related phenotypes and promotes the development of metabolic abnormalities. Am J Pathol, 180(3):963–972, Mar 2012.

32 K. Ozaki, H. Sato, A. Iida, H. Mizuno, T. Nakamura, Y. Miyamoto, A. Takahashi, T. Tsunoda, S. Ikegawa, N. Kamatani, M. Hori, Y. Nakamura, and T. Tanaka. A functional SNP in PSMA6 confers risk of myocardial infarction in the Japanese population. Nat Genet, 38(8):921–925, Aug 2006.

33 M. G. Heckman, A. I. Soto-Ortolaza, N. N. Diehl, S. Rayaprolu, T. G. Brott, Z. K. Wszolek, J. F. Meschia, and O. A. Ross. Genetic variants associated with myocardial infarction in the PSMA6 gene and Chr9p21 are also associated with ischaemic stroke. Eur J Neurol, 20(2):300–308, Feb 2013.

34 T. Sjakste, M. Kalis, I. Poudziunas, V. Pirags, M. Lazdins, L. Groop, and N. Sjakste. Association of microsatellite polymorphisms of the human 14q13.2 region with type 2 diabetes mellitus in Latvian and Finnish populations. Ann Hum Genet, 71(Pt 6):772–776, Nov 2007.

35 S. S. Wing. The UPS in diabetes and obesity. BMC Biochem, 9 Suppl 1:S6, Oct 2008.

36 E. L. Webb, M. F. Rudd, G. S. Sellick, R. El Galta, L. Bethke, W. Wood, O. Fletcher, S. Penegar, L. Withey, M. Qureshi, N. Johnson, I. Tomlinson, R. Gray, J. Peto, and R. S. Houlston. Search for low penetrance alleles for colorectal cancer through a scan of 1467 non-synonymous SNPs in 2575 cases and 2707 controls with validation by kin-cohort analysis of 14 704 first-degree relatives. Hum Mol Genet, 15(21):3263–3271, Nov 2006.

37 C. Zeng, K. Matsuda, W. H. Jia, J. Chang, S. S. Kweon, Y. B. Xiang, A. Shin, S. H. Jee, D. H. Kim, B. Zhang, Q. Cai, X. Guo, J. Long, N. Wang, R. Courtney, Z. Z. Pan, C. Wu, A. Takahashi, M. H. Shin, K. Matsuo, F. Matsuda, Y. T. Gao, J. H. Oh, S. Kim, K. J. Jung, Y. O. Ahn, Z. Ren, H. L. Li, J. Wu, J. Shi, W. Wen, G. Yang, B. Li, B. T. Ji, H. Brenner, R. E. Schoen, S. Küry, S. B. Gruber, F. R. Schumacher, S. L. Stenzel, G. Casey, J. L. Hopper, M. A. Jenkins, H. R. Kim, J. Y. Jeong, J. W. Park, K. Tajima, S. H. Cho, M. Kubo, X. O. Shu, D. Lin, Y. X. Zeng, W. Zheng, J. A. Baron, S. I. Berndt, S. Bezieau, H. Brenner, B. J. Caan, C. S. Carlson, G. Casey, A. T. Chan, J. Chang-Claude, S. J. Chanock, D. V. Conti, K. Curtis, D. Duggan, C. S. Fuchs, S. Gallinger, E. L. Giovannucci, S. B. Gruber, R. W. Haile, T. A. Harrison, R. B. Hayes, M. Hoffmeister, J. L. Hopper, L. Hsu, T. J. Hudson, D. J. Hunter, C. M. Hutter, R. D. Jackson, M. A. Jenkins, S. Jiao, S. Küry, L. Le Marchand, M. Lemire, N. M. Lindor, J. Ma, P. A. Newcomb, U. Peters, J. D. Potter, C. Qu, R. E. Schoen, F. R. Schumacher, D. Seminara, M. L. Slattery, S. N. Thibodeau, E. White, B. W. Zanke, K. Blalock, P. T. Campbell, G. Casey, D. V. Conti, C. K. Edlund, J. Figueiredo, W. J. Gauderman, J. Gong, R. C. Green, S. B. Gruber, J. F. Harju, T. A. Harrison, E. J. Jacobs, M. A. Jenkins, S. Jiao, L. Li, Y. Lin, F. J. Manion, V. Moreno, B. Mukherjee, U. Peters, L. Raskin, F. R. Schumacher, D. Seminara, G. Severi, S. L. Stenzel, and D. C. Thomas. Identification of Susceptibility Loci and Genes for Colorectal Cancer Risk. Gastroenterology, 150(7):1633–1645, 06 2016.

38 K. Shameer, J. C. Denny, K. Ding, H. Jouni, D. R. Crosslin, M. de Andrade, C. G. Chute, P. Peissig, J. A. Pacheco, R. Li, L. Bastarache, A. N. Kho, M. D. Ritchie, D. R. Masys, R. L. Chisholm, E. B. Larson, C. A. McCarty, D. M. Roden, G. P. Jarvik, and I. J. Kullo. A genome- and phenome-wide association study to identify genetic variants influencing platelet count and volume and their pleiotropic effects. Hum Genet, 133(1):95–109, Jan 2014.

39 P. E. Stuart, R. P. Nair, E. Ellinghaus, J. Ding, T. Tejasvi, J. E. Gudjonsson, Y. Li, S. Weidinger, B. Eberlein, C. Gieger, H. E. Wichmann, M. Kunz, R. Ike, G. G. Krueger, A. M. Bowcock, U. Mrowietz, H. W. Lim, J. J. Voorhees, G. R. Abecasis, M. Weichenthal, A. Franke, P. Rahman, D. D. Gladman, and J. T. Elder. Genome-wide association analysis identifies three psoriasis susceptibility loci. Nat Genet, 42(11):1000–1004, Nov 2010.

40 E. Iio, K. Matsuura, N. Nishida, S. Maekawa, N. Enomoto, M. Nakagawa, N. Sakamoto, H. Yatsuhashi, M. Kurosaki, N. Izumi, Y. Hiasa, N. Masaki, T. Ide, K. Hino, A. Tamori, M. Honda, S. Kaneko, S. Mochida, H. Nomura, S. Nishiguchi, C. Okuse, Y. Itoh, H. Yoshiji, I. Sakaida, K. Yamamoto, H. Watanabe, S. Hige, A. Matsumoto, E. Tanaka, K. Tokunaga, and Y. Tanaka. Genome-wide association study identifies a PSMD3 variant associated with neutropenia in interferon-based therapy for chronic hepatitis C. Hum Genet, 134(3):279–289, Mar 2015.

41 X. Song, G. Zhu, S. Hou, Y. Ren, M. W. Amjid, W. Li, and W. Guo.). Front Plant Sci, 12:695503, 2021.

42 A. Belle, A. Tanay, L. Bitincka, R. Shamir, and E. K. O’Shea. Quantification of protein half-lives in the budding yeast proteome. Proc Natl Acad Sci U S A, 103(35):13004–13009, Aug 2006.

43 R. Christiano, N. Nagaraj, F. Frohlich, and T. C. Walther. Global proteome turnover analyses of the Yeasts S. cerevisiae and S. pombe. Cell Rep, 9(5):1959–1965, Dec 2014.

44 B. Schwanhausser, D. Busse, N. Li, G. Dittmar, J. Schuchhardt, J. Wolf, W. Chen, and M. Selbach. Global quantification of mammalian gene expression control. Nature, 473(7347):337–342, May 2011.

45 D. A. Kraut and A. Matouschek. Proteasomal degradation from internal sites favors partial proteolysis via remote domain stabilization. ACS Chem Biol, 6(10):1087–1095, Oct 2011.

46 S. Prakash, L. Tian, K. S. Ratliff, R. E. Lehotzky, and A. Matouschek. An unstructured initiation site is required for efficient proteasome-mediated degradation. Nat Struct Mol Biol, 11(9):830– 837, Sep 2004.

47 K. Martinez-Fonts, C. Davis, T. Tomita, S. Elsasser, A. R. Nager, Y. Shi, D. Finley, and A. Matouschek. The proteasome 19S cap and its ubiquitin receptors provide a versatile recognition platform for substrates. Nat Commun, 11(1):477, 01 2020.

48 D. A. Kraut, E. Israeli, E. K. Schrader, A. Patil, K. Nakai, D. Nanavati, T. Inobe, and A. Matouschek. Sequence-and species-dependence of proteasomal processivity. ACS Chem Biol, 7(8):1444–1453, Aug 2012.

49 H. Yu, A. K. Singh Gautam, S. R. Wilmington, D. Wylie, K. Martinez-Fonts, G. Kago, M. Warburton, S. Chavali, T. Inobe, I. J. Finkelstein, M. M. Babu, and A. Matouschek. Conserved Sequence Preferences Contribute to Substrate Recognition by the Proteasome. J Biol Chem, 291(28):14526–14539, Jul 2016.

50 M. A. Hoyt, J. Zich, J. Takeuchi, M. Zhang, C. Govaerts, and P. Coffino. Glycine-alanine repeats impair proper substrate unfolding by the proteasome. EMBO J, 25(8):1720–1729, Apr 2006.

51 P. Koodathingal, N. E. Jaffe, D. A. Kraut, S. Prakash, S. Fishbain, C. Herman, and A. Matouschek. ATP-dependent proteases differ substantially in their ability to unfold globular proteins. J Biol Chem, 284(28):18674–18684, Jul 2009.

52 J. Abi Habib, E. De Plaen, V. Stroobant, D. Zivkovic, M. P. Bousquet, B. Guillaume, K. Wahni, J. Messens, A. Busse, N. Vigneron, and B. J. Van den Eynde. Efficiency of the four proteasome subtypes to degrade ubiquitinated or oxidized proteins. Sci Rep, 10(1):15765, 09 2020.

53 J. Abi Habib, J. Lesenfants, N. Vigneron, and B. J. Van den Eynde. Functional Differences between Proteasome Subtypes. Cells, 11(3), 01 2022.

54 M. Bajorek, D. Finley, and M. H. Glickman. Proteasome disassembly and downregulation is correlated with viability during stationary phase. Curr Biol, 13(13):1140–1144, Jul 2003.

55 T. Mayor, M. Sharon, and M. H. Glickman. Tuning the proteasome to brighten the end of the journey. Am J Physiol Cell Physiol, 311(5):C793–C804, Nov 2016.

56 R. Shringarpure, T. Grune, J. Mehlhase, and K. J. Davies. Ubiquitin conjugation is not required for the degradation of oxidized proteins by proteasome. J Biol Chem, 278(1):311–318, Jan 2003.

57 J. S. Bloom, I. M. Ehrenreich, W. T. Loo, T. L. Lite, and L. Kruglyak. Finding the sources of missing heritability in a yeast cross. Nature, 494(7436):234–237, Feb 2013.

58 H. C. Yen, Q. Xu, D. M. Chou, Z. Zhao, and S. J. Elledge. Global protein stability profiling in mammalian cells. Science, 322(5903):918–923, Nov 2008.

59 Y. Geffen, A. Appleboim, R. G. Gardner, N. Friedman, R. Sadeh, and T. Ravid. Mapping the Landscape of a Eukaryotic Degronome. Mol Cell, 63(6):1055–1065, 09 2016.

60 H. Ella, Y. Reiss, and T. Ravid. The Hunt for Degrons of the 26S Proteasome. Biomolecules, 9(6), 06 2019.

61 M. Zhang, C. M. Pickart, and P. Coffino. Determinants of proteasome recognition of ornithine decarboxylase, a ubiquitin-independent substrate. EMBO J, 22(7):1488–1496, Apr 2003.

62 J. Takeuchi, H. Chen, M. A. Hoyt, and P. Coffino. Structural elements of the ubiquitin-independent proteasome degron of ornithine decarboxylase. Biochem J, 410(2):401–407, Mar 2008.

63 Y. Xie and A. Varshavsky. RPN4 is a ligand, substrate, and transcriptional regulator of the 26S proteasome: a negative feedback circuit. Proc. Natl. Acad. Sci. U.S.A., 98(6):3056–3061, Mar 2001.

64 S. W. Ha, D. Ju, and Y. Xie. The N-terminal domain of Rpn4 serves as a portable ubiquitin-independent degron and is recognized by specific 19S RP subunits. Biochem Biophys Res Commun, 419(2):226–231, Mar 2012.

65 D. Ju and Y. Xie. Proteasomal degradation of RPN4 via two distinct mechanisms, ubiquitin-dependent and -independent. J Biol Chem, 279(23):23851–23854, Jun 2004.

66 J. Erales and P. Coffino. Ubiquitin-independent proteasomal degradation. Biochim Biophys Acta, 1843(1):216–221, Jan 2014.

67 M. A. Hoyt, M. Zhang, and P. Coffino. Ubiquitin-independent mechanisms of mouse ornithine decarboxylase degradation are conserved between mammalian and fungal cells. J Biol Chem, 278(14):12135–12143, Apr 2003.

68 G. Mannhaupt, R. Schnall, V. Karpov, I. Vetter, and H. Feldmann. Rpn4p acts as a transcription factor by binding to PACE, a nonamer box found upstream of 26S proteasomal and other genes in yeast. FEBS Lett, 450(1-2):27–34, Apr 1999.

69 A. V. Morozov, D. S. Spasskaya, D. S. Karpov, and V. L. Karpov. The central domain of yeast transcription factor Rpn4 facilitates degradation of reporter protein in human cells. FEBS Lett, 588(20):3713–3719, Oct 2014.

70 M. A. Hoyt, M. Zhang, and P. Coffino. Probing the ubiquitin/proteasome system with ornithine decarboxylase, a ubiquitin-independent substrate. Methods Enzymol, 398:399–413, 2005.

71 I. Momose, D. Tatsuda, S. Ohba, T. Masuda, D. Ikeda, and A. Nomoto. In vivo imaging of proteasome inhibition using a proteasome-sensitive fluorescent reporter. Cancer Sci, 103(9):1730– 1736, Sep 2012.

72 A. Khmelinskii and M. Knop. Analysis of protein dynamics with tandem fluorescent protein timers. Methods Mol. Biol., 1174:195–210, 2014.

73 A. Khmelinskii, M. Meurer, C. T. Ho, B. Besenbeck, J. F?ller, M. K. Lemberg, B. Bukau, A. Mogk, and M. Knop. Incomplete proteasomal degradation of green fluorescent proteins in the context of tandem fluorescent protein timers. Mol. Biol. Cell, 27(2):360–370, Jan 2016.

74 J. D. Pedelacq, S. Cabantous, T. Tran, T. C. Terwilliger, and G. S. Waldo. Engineering and characterization of a superfolder green fluorescent protein. Nat. Biotechnol., 24(1):79–88, Jan 2006.

75 N. C. Shaner, R. E. Campbell, P. A. Steinbach, B. N. Giepmans, A. E. Palmer, and R. Y. Tsien. Improved monomeric red, orange and yellow fluorescent proteins derived from Discosoma sp. red fluorescent protein. Nat. Biotechnol., 22(12):1567–1572, Dec 2004.

76 A. Khmelinskii, P. J. Keller, A. Bartosik, M. Meurer, J. D. Barry, B. R. Mardin, A. Kaufmann, S. Trautmann, M. Wachsmuth, G. Pereira, W. Huber, E. Schiebel, and M. Knop. Tandem fluorescent protein timers for in vivo analysis of protein dynamics. Nat. Biotechnol., 30(7):708– 714, Jun 2012.

77 F. W. Albert, J. S. Bloom, J. Siegel, L. Day, and L. Kruglyak. Genetics of trans-regulatory variation in gene expression. Elife, 7, 07 2018.

78 F. W. Albert, S. Treusch, A. H. Shockley, J. S. Bloom, and L. Kruglyak. Genetics of single-cell protein abundance variation in large yeast populations. Nature, 506(7489):494–497, Feb 2014.

79 C. Brion, S. M. Lutz, and F. W. Albert. Simultaneous quantification of mRNA and protein in single cells reveals post-transcriptional effects of genetic variation. Elife, 9, Nov 2020.

80 M. A. Hoyt, S. McDonough, S. A. Pimpl, H. Scheel, K. Hofmann, and P. Coffino. A genetic screen for Saccharomyces cerevisiae mutants affecting proteasome function, using a ubiquitin-independent substrate. Yeast, 25(3):199–217, Mar 2008.

81 J. Takeuchi, H. Chen, and P. Coffino. Proteasome substrate degradation requires association plus extended peptide. EMBO J, 26(1):123–131, Jan 2007.

82 D. Ju, H. Xu, X. Wang, and Y. Xie. The transcription activation domain of Rpn4 is separate from its degrons. Int J Biochem Cell Biol, 42(2):282–286, Feb 2010.

83 I. M. Ehrenreich, J. P. Gerke, and L. Kruglyak. Genetic dissection of complex traits in yeast: insights from studies of gene expression and other phenotypes in the BYxRM cross. Cold Spring Harb Symp Quant Biol, 74:145–153, 2009.

84 I. M. Ehrenreich, N. Torabi, Y. Jia, J. Kent, S. Martis, J. A. Shapiro, D. Gresham, A. A. Caudy, and L. Kruglyak. Dissection of genetically complex traits with extremely large pools of yeast segregants. Nature, 464(7291):1039–1042, Apr 2010.

85 R. W. Michelmore, I. Paran, and R. V. Kesseli. Identification of markers linked to disease-resistance genes by bulked segregant analysis: a rapid method to detect markers in specific genomic regions by using segregating populations. Proc Natl Acad Sci U S A, 88(21):9828– 9832, Nov 1991.

86 A. Varshavsky. The N-end rule pathway and regulation by proteolysis. Protein Sci., 20(8):1298– 1345, Aug 2011.

87 C. S. Hwang, A. Shemorry, and A. Varshavsky. N-terminal acetylation of cellular proteins creates specific degradation signals. Science, 327(5968):973–977, Feb 2010.

88 A. Bachmair, D. Finley, and A. Varshavsky. In vivo half-life of a protein is a function of its amino-terminal residue. Science, 234(4773):179–186, Oct 1986.

89 R. T. Baker and A. Varshavsky. Yeast N-terminal amidase. A new enzyme and component of the N-end rule pathway. J Biol Chem, 270(20):12065–12074, May 1995.

90 R. B. Wickner. MKT1, a nonessential Saccharomyces cerevisiae gene with a temperature-dependent effect on replication of M2 double-stranded RNA. J Bacteriol, 169(11):4941–4945, Nov 1987.

91 T. Tadauchi, T. Inada, K. Matsumoto, and K. Irie. Posttranscriptional regulation of HO expression by the Mkt1-Pbp1 complex. Mol Cell Biol, 24(9):3670–3681, May 2004.

92 H. Sinha, B. P. Nicholson, L. M. Steinmetz, and J. H. McCusker. Complex genetic interactions in a quantitative trait locus. PLoS Genet, 2(2):e13, Feb 2006.

93 A. M. Deutschbauer and R. W. Davis. Quantitative trait loci mapped to single-nucleotide resolution in yeast. Nat Genet, 37(12):1333–1340, Dec 2005.

94 K. Tanaka, M. Nakafuku, F. Tamanoi, Y. Kaziro, K. Matsumoto, and A. Toh-e. IRA2, a second gene of Saccharomyces cerevisiae that encodes a protein with a domain homologous to mammalian ras GTPase-activating protein. Mol Cell Biol, 10(8):4303–4313, Aug 1990.

95 E. N. Smith and L. Kruglyak. Gene-environment interaction in yeast gene expression. PLoS Biol, 6(4):e83, Apr 2008.

96 S. Lutz, K. Van Dyke, M. A. Feraru, and F. W. Albert. Multiple epistatic DNA variants in a single gene affect gene expression in trans. Genetics, 220(1), 01 2022.

97 L. A. Howell, A. K. Peterson, and R. J. Tomko. Proteasome subunit a1 overexpression preferentially drives canonical proteasome biogenesis and enhances stress tolerance in yeast. Sci Rep, 9(1):12418, 08 2019.

98 D. Vilchez, L. Boyer, I. Morantte, M. Lutz, C. Merkwirth, D. Joyce, B. Spencer, L. Page, E. Masliah, W. T. Berggren, F. H. Gage, and A. Dillin. Increased proteasome activity in human embryonic stem cells is regulated by PSMD11. Nature, 489(7415):304–308, Sep 2012.

99 C. G. de Boer and T. R. Hughes. YeTFaSCo: a database of evaluated yeast transcription factor sequence specificities. Nucleic Acids Res, 40(Database issue):D169–179, Jan 2012.

100 G. Owsianik, L. Balzi l, and M. Ghislain. Control of 26S proteasome expression by transcription factors regulating multidrug resistance in Saccharomyces cerevisiae. Mol Microbiol, 43(5):1295–1308, Mar 2002.

101 X. Wang, J. Yen, P. Kaiser, and L. Huang. Regulation of the 26S proteasome complex during oxidative stress. Sci Signal, 3(151):ra88, Dec 2010.

102 H. Salin, V. Fardeau, E. Piccini, G. Lelandais, V. Tanty, S. Lemoine, C. Jacq, and F. Devaux. Structure and properties of transcriptional networks driving selenite stress response in yeasts. BMC Genomics, 9:333, Jul 2008.

103 J. Peter, M. De Chiara, A. Friedrich, J. X. Yue, D. Pflieger, A. Bergström, A. Sigwalt, B. Barre, K. Freel, A. Llored, C. Cruaud, K. Labadie, J. M. Aury, B. Istace, K. Lebrigand, P. Barbry, S. Engelen, A. Lemainque, P. Wincker, G. Liti, and J. Schacherer. Genome evolution across 1,011 Saccharomyces cerevisiae isolates. Nature, 556(7701):339–344, 04 2018.

104 A. N. Zyrina, E. A. Smirnova, O. V. Markova, F. F. Severin, and D. A. Knorre. Mitochondrial Superoxide Dismutase and Yap1p Act as a Signaling Module Contributing to Ethanol Tolerance of the Yeast Saccharomyces cerevisiae. Appl Environ Microbiol, 83(3), 02 2017.

105 K. Flick and P. Kaiser. Protein degradation and the stress response. Semin Cell Dev Biol, 23(5):515–522, Jul 2012.

106 J. Hanna, A. Meides, D. P. Zhang, and D. Finley. A ubiquitin stress response induces altered proteasome composition. Cell, 129(4):747–759, May 2007.

107 A. V. Gomes. Genetics of proteasome diseases. Scientifica (Cairo), 2013:637629, 2013.

108 K. Arima, A. Kinoshita, H. Mishima, N. Kanazawa, T. Kaneko, T. Mizushima, K. Ichinose, H. Nakamura, A. Tsujino, A. Kawakami, M. Matsunaka, S. Kasagi, S. Kawano, S. Kumagai, K. Ohmura, T. Mimori, M. Hirano, S. Ueno, K. Tanaka, M. Tanaka, I. Toyoshima, H. Sugino, A. Yamakawa, K. Tanaka, N. Niikawa, F. Furukawa, S. Murata, K. Eguchi, H. Ida, and K. Yoshiura. Proteasome assembly defect due to a proteasome subunit beta type 8 (PSMB8) mutation causes the autoinflammatory disorder, Nakajo-Nishimura syndrome. Proc Natl Acad Sci U S A, 108(36):14914–14919, Sep 2011.

109 D. Komander and M. Rape. The ubiquitin code. Annu Rev Biochem, 81:203–229, 2012.

110 F. Ohtake, Y. Saeki, S. Ishido, J. Kanno, and K. Tanaka. The K48-K63 Branched Ubiquitin Chain Regulates NF-KB Signaling. Mol Cell, 64(2):251–266, 10 2016.

111 M. E. French, C. F. Koehler, and T. Hunter. Emerging functions of branched ubiquitin chains. Cell Discov, 7(1):6, Jan 2021.

112 Y. Murakami, S. Matsufuji, T. Kameji, S. Hayashi, K. Igarashi, T. Tamura, K. Tanaka, and A. Ichihara. Ornithine decarboxylase is degraded by the 26S proteasome without ubiquitination. Nature, 360(6404):597–599, Dec 1992.

113 S. Kors, K. Geijtenbeek, E. Reits, and S. Schipper-Krom. Regulation of Proteasome Activity by (Post-)transcriptional Mechanisms. Front Mol Biosci, 6:48, 2019.

114 C. Pla-Prats and N. H. Thoma. Quality control of protein complex assembly by the ubiquitin-proteasome system. Trends Cell Biol, Mar 2022.

115 C. Padovani, P. Jevtic, and M. Rape. Quality control of protein complex composition. Mol Cell, 82(8):1439–1450, Apr 2022.

116 Y. Saeki, A. Toh-E, T. Kudo, H. Kawamura, and K. Tanaka. Multiple proteasome-interacting proteins assist the assembly of the yeast 19S regulatory particle. Cell, 137(5):900–913, May 2009.

117 A. Khmelinskii, E. Blaszczak, M. Pantazopoulou, B. Fischer, D. J. Omnus, G. Le Dez, A. Brossard, A. Gunnarsson, J. D. Barry, M. Meurer, D. Kirrmaier, C. Boone, W. Huber, G. Rabut, P. O. Ljungdahl, and M. Knop. Protein quality control at the inner nuclear membrane. Nature, 516(7531):410–413, Dec 2014.

118 M. Zhang, A. I. MacDonald, M. A. Hoyt, and P. Coffino. Proteasomes begin ornithine decarboxylase digestion at the C terminus. J Biol Chem, 279(20):20959–20965, May 2004.

119 A. L. Goldstein and J. H. McCusker. Three new dominant drug resistance cassettes for gene disruption in Saccharomyces cerevisiae. Yeast, 15(14):1541–1553, Oct 1999.

120 J. H. Zwolshen and J. K. Bhattacharjee. Genetic and biochemical properties of thialysine-resistant mutants of Saccharomyces cerevisiae. J Gen Microbiol, 122(2):281–287, Feb 1981.

121 A. Baryshnikova, M. Costanzo, S. Dixon, F. J. Vizeacoumar, C. L. Myers, B. Andrews, and C. Boone. Synthetic genetic array (SGA) analysis in Saccharomyces cerevisiae and Schizosac-charomyces pombe. Methods Enzymol, 470:145–179, 2010.

122 E. Kuzmin, M. Costanzo, B. Andrews, and C. Boone. Synthetic Genetic Array Analysis. Cold Spring Harb Protoc, 2016(4):pdb.prot088807, Apr 2016.

123 R. D. Gietz and R. H. Schiestl. High-efficiency yeast transformation using the LiAc/SS carrier DNA/PEG method. Nat Protoc, 2(1):31–34, 2007.

124 A. C. Ward. Rapid analysis of yeast transformants using colony-PCR. Biotechniques, 13(3):350, Sep 1992.

125 R Core Team. R: A Language and Environment for Statistical Computing. R Foundation for Statistical Computing, Vienna, Austria, 2021.

126 F. Hahne, N. LeMeur, R. R. Brinkman, B. Ellis, P. Haaland, D. Sarkar, J. Spidlen, E. Strain, and R. Gentleman. flowCore: a Bioconductor package for high throughput flow cytometry. BMC Bioinformatics, 10:106, Apr 2009.

127 Douglas Bates, Martin Machler, Ben Bolker, and Steve Walker. Fitting linear mixed-effects models using lme4. Journal of Statistical Software, 67(1):1–48, 2015.

128 Alexandra Kuznetsova, Per B. Brockhoff, and Rune H. B. Christensen. lmerTest package: Tests in linear mixed effects models. Journal of Statistical Software, 82(13):1–26, 2017.

129 H. Li and R. Durbin. Fast and accurate short read alignment with Burrows-Wheeler transform. Bioinformatics, 25(14):1754–1760, Jul 2009.

130 H. Li, B. Handsaker, A. Wysoker, T. Fennell, J. Ruan, N. Homer, G. Marth, G. Abecasis, and R. Durbin. The Sequence Alignment/Map format and SAMtools. Bioinformatics, 25(16):2078– 2079, Aug 2009.

131 M. D. Edwards and D. K. Gifford. High-resolution genetic mapping with pooled sequencing. BMC Bioinformatics, 13 Suppl 6:S8, Apr 2012.

132 W. J. Kent, C. W. Sugnet, T. S. Furey, K. M. Roskin, T. H. Pringle, A. M. Zahler, and D. Haussler. The human genome browser at UCSC. Genome Res, 12(6):996–1006, Jun 2002.

133 M. Haeussler, K. Schönig, H. Eckert, A. Eschstruth, J. Mianné, J. B. Renaud, S. Schneider-Maunoury, A. Shkumatava, L. Teboul, J. Kent, J. S. Joly, and J. P. Concordet. Evaluation of off-target and on-target scoring algorithms and integration into the guide RNA selection tool CRISPOR. Genome Biol, 17(1):148, 07 2016.

134 J. G. Doench, N. Fusi, M. Sullender, M. Hegde, E. W. Vaimberg, K. F. Donovan, I. Smith, Z. Tothova, C. Wilen, R. Orchard, H. W. Virgin, J. Listgarten, and D. E. Root. Optimized sgRNA design to maximize activity and minimize off-target effects of CRISPR-Cas9. Nat Biotechnol, 34(2):184–191, Feb 2016.

135 C. S. Hoffman and F. Winston. A ten-minute DNA preparation from yeast efficiently releases autonomous plasmids for transformation of Escherichia coli. Gene, 57(2-3):267–272, 1987.

136 R. M. Horton, H. D. Hunt, S. N. Ho, J. K. Pullen, and L. R. Pease. Engineering hybrid genes without the use of restriction enzymes: gene splicing by overlap extension. Gene, 77(1):61–68, Apr 1989.

